# A Mechanistic Theory Unifying Value and Schema in the Prefrontal Cortex

**DOI:** 10.64898/2026.10.17.759791

**Authors:** Joseph Warren, Kristopher T. Jensen, Timothy E. J. Behrens, James C. R. Whittington

**Affiliations:** Sainsbury Wellcome Centre, University College London, London, UK; Oxford Centre for Integrative Neuroimaging, University of Oxford, Oxford, UK; Department of Experimental Psychology, University of Oxford, UK

## Abstract

Animals exhibit behavioural flexibility by making reward-optimal decisions in unfamiliar situations. Computing value from minimal reward feedback requires leveraging known structure in the world. The Prefrontal Cortex (PFC) plays a pivotal role in both value computation and learning world structure, however mechanistic explanations for each—within the frameworks of Reinforcement Learning and Cognitive Maps—are largely distinct. We develop a mechanistic model of PFC that unifies value and structure through behavioural flexibility. In this model, value acts as a control signal that gates structured working memory representations of the current context in order to ultimately control behaviour. This is a significant departure from the conventional perspective of value, in which it is used for action readout. Critically, this model generalises to novel contexts without the need for additional synaptic plasticity. We demonstrate that recurrent neural networks (RNNs) meta-trained with either reinforcement learning or supervised learning recapitulate this exact model on various value-based tasks. These results not only provide a parsimonious explanation of both the value and schema frameworks of PFC function, but also propose a new mechanistic interpretation of the role of PFC in flexible decision making.

## 1 Introduction

Animals are capable of flexibly adapting behaviour to new contexts in order to obtain reward. While such behavioural adaptation requires learning from reward feedback, this feedback is often extremely minimal, which is at odds with rote trial-and-error learning prescribed by conventional reinforcement learning (RL). Instead, this learning is more analogous to inference, with optimal behaviours deduced by leveraging pre-learned structures from different but related experiences. Pertinently, learning from reward and representing structure (cognitive maps) are two well established functions of PFC, but we do not have a mechanistic understanding of how they unite to support flexible decision-making.

Value signals are ubiquitous in PFC (Figure 1a) and are causally involved in decision making (Knudsen & Wallis, 2022; Ballesta et al., 2020; Gardner et al., 2020; Camille et al., 2011), however there is no clear framework for why they look the way they do. For instance, individual PFC neurons exhibit tunings that appear to be a computational substrate for decision-making, such as expected value, reward magnitude, reward probability, and cost (Padoa-Schioppa & Assad, 2006, 2008; Kennerley et al., 2009, 2011). Some of these responses represent the value of each possible action, which would provide a natural substrate for decision-making. However, a large proportion of responses instead represent value in chosen (Padoa-Schioppa & Assad, 2006) and unchosen (Boorman et al., 2011) coordinates—or similarly in best, middle and worst coordinates (Knudsen & Wallis, 2020) for three options—irrespective of the actual action identity (Padoa-Schioppa & Assad, 2008). It is not clear why representing value in an action-invariant best-to-worst frame of reference would be useful for controlling behaviour, since it does not easily allow actions to be directly read-out, unlike conventional RL.

**Figure 1:**
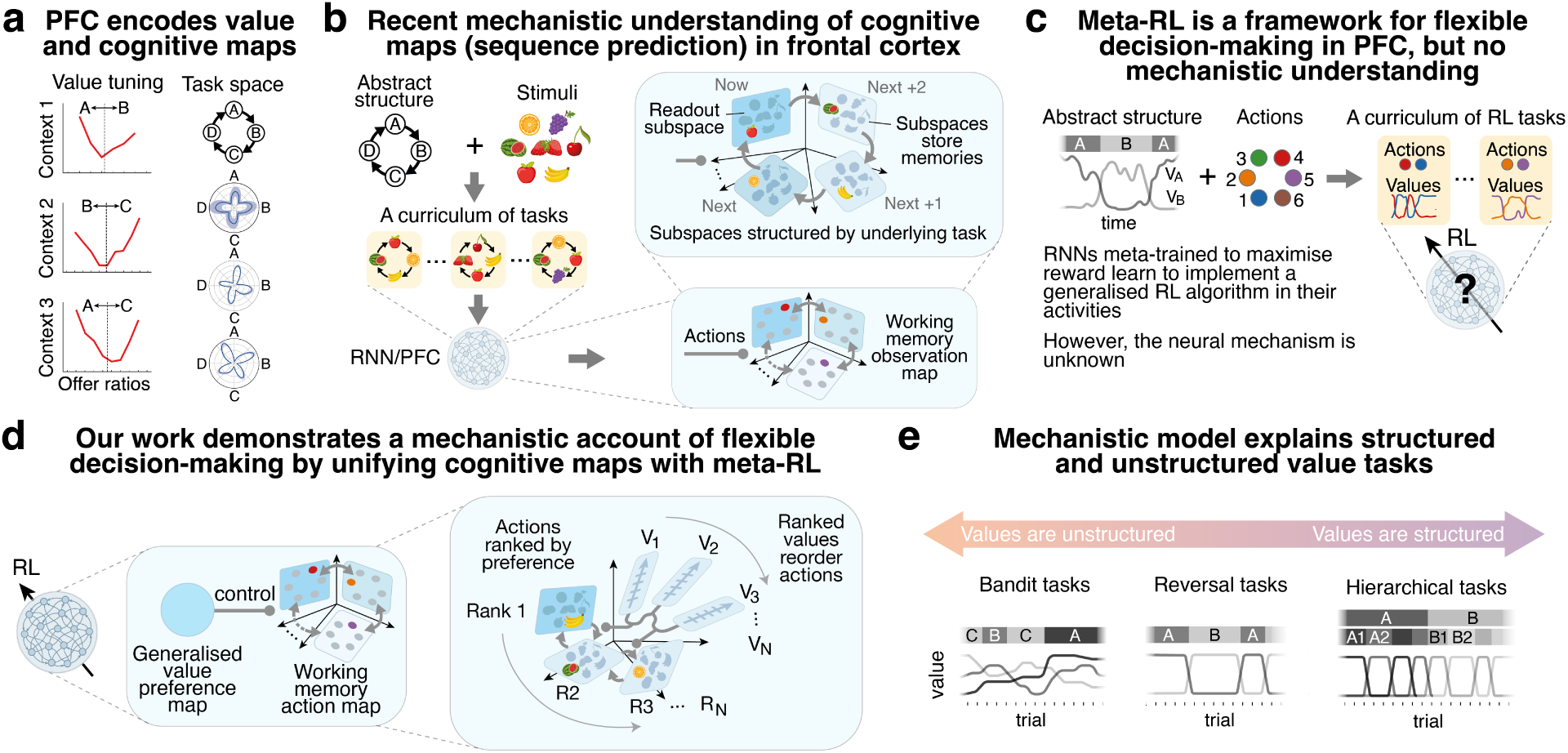
Mechanistic unification of value and sequence working memory (cognitive maps) in Prefrontal Cortex. **a)** Left: an orbitofrontal (OFC) neuron tuned to chosen (relative) value which is invariant both to the identity of the better option (U-shape) and to which options are available from juice A, B, or C. Adapted from Padoa-Schioppa & Assad (2008). Right: three VmPFC neurons tuned to task space on a 4-goal navigation task, adapted from El-Gaby et al. (2024). **b)** Learning a map can be framed as a sequence memory task—predicting what will be seen next—in which each problem is a combination of an abstract structure (e.g., a 4-loop) and arbitrary sensory stimuli (top). Meta-trained RNNs and PFC generalise to unseen problems and both learn the same solution (bottom) in which memories of the problem’s stimuli are stored in the neural activity of subspaces structured according to the underlying task structure. Movement within the problem (e.g., external actions clockwise / counter-clockwise) induces a corresponding movement of the memories between subspaces, to ensure the right memory can be readout from the readout subspace. **c)** Meta-RL-trained RNNs demonstrate similar ability *in the decision-making domain* by generalising to new problems with familiar structure. Here an example meta-RL task—generalised reversal learning—is shown in which the values of the two options are anti-correlated, but where each problem has different options. **d)** Our proposed model mechanistically unifies value with structured working memory. Structured subspaces store the ‘available’ options in the context/problem, just as stimuli were stored for the cognitive map. Internal value signals control the re-ordering of memories between the subspaces, just as external actions did for cognitive maps. Subspaces and values are organised by ranked preference of the *N* relevant options in the current context. Comparison between the ranked values acts as a control signal for swaps of specific subspace contents. **e)** Proposed model applies to unstructured and structured value-based tasks. Structure refers to the dependence of action values on one another.

Intriguingly a parallel literature of PFC in sequential prediction tasks shows a similar relative ordering of represented variables. In such tasks, subjects are shown a sequence of items and are required to predict the subsequent item(s). Critically, these sequences are *structured*—they are governed by a common set of transformations that operate on past items to determine the next item in the sequence (Figure 1b shows an example ‘loop’ sequence structure with action-dependent transformations). Accurate prediction requires learning these transformations, and this is done slowly after experiencing many example sequences. Once learnt, the transformations can be applied instantaneously to new example sequences, allowing rapid inferences on new data—a form of learning known in AI as meta-learning. Models and data (Whittington et al., 2025; El-Gaby et al., 2024) suggest a circuit mechanism for these rapid inferences in PFC: the items to be remembered in a sequence are stored in separate working memory (WM) subspaces (‘slots’), and these subspaces are connected to one another with connectivity that embeds the transformations (task structure), so that the memory contents can be moved between subspaces. This memory shuffling process means that 1) all the memories for a given sequence are stored without interfering with each other and 2) that the right memory occupies the read-out subspace at the right time. In other words, the connectivity between subspaces—*meta-learned* to embed the underlying sequence structure—order the memories into relative coordinates depending on when they will likely be read out.

Many value and reinforcement learning tasks also involve a training period where an animal learns the task structure. Furthermore, although such studies are less common, when tested, animals will typically generalise this task structure seamlessly to new spatial layouts (Samborska et al., 2021; El-Gaby et al., 2024), objects (Harlow, 1949) or sensory stimuli (Slotnick et al., 2000; Zhou et al., 2021; Hosokawa et al., 2018). Hence meta-learning provides an attractive framework for understanding value-guided decision making tasks.

Indeed, meta-reinforcement learning (meta-RL) has recently been proposed to unify value and structure in PFC (meta-RL; Figure 1c) (Wang et al., 2018). The key insight is that the dynamics of PFC can learn to effectively implement their own RL algorithm, different from the RL algorithm that is used to train them. This learnt RL algorithm can be extremely sophisticated, performing task-specific inferences from sparse rewards that allow rapid and flexible behaviours online. This learned RL algorithm requires no synaptic change, only changes in activity; its RL is just like *inference*. For example, in the classic Harlow learning set task (Harlow, 1949), being unrewarded on one item immediately tells the subject the other item is rewarded, without ever needing to sample it. Such inferences are much easier with an RL algorithm encoded in dynamics (rather than synapses in conventional RL) since values can be encoded in neural activity and their relationships in synaptic connections meaning an update to one value can cause an update to all the others.

The meta-learning account provides a natural explanation for flexible behaviour, and for why value and structured representations are found in PFC neurons (Wang et al., 2018). However, it is unknown what is the mechanistic circuit implementation of such meta-learnt RL algorithms. We therefore do not have a clear picture of why some representations emerge and not others, nor how different parts of the representational space interact to solve a new task. Unlike in the sequence prediction (working memory) tasks above, we do not understand the circuit mechanism, making it difficult to have clear predictions for neural data.

In this paper we propose a mechanistic theory that unifies PFC value and structured representations, and we validate this theory *in silico* by meta-training recurrent networks on a variety of reward-based decision making tasks. We demonstrate that these networks lean to encode action-invariant values in their activity, and this value code serves as a control signal that reorders working memory representations of action preferences to guide choices. In doing so, we show that the RL algorithms learnt by meta-learners are closely related to the solutions learnt for structured sequence prediction.

## 2 Model: structured representations for value-based decision-making

To understand representations of task structure in RL, we consider the meta-learning setting, where networks are trained to embed structure common across contexts. Here, a task is defined as a set of problems (contexts) that share the same underlying value structure, but have different relevant actions or map states to different stimuli (Figure 1c).

We train RNNs with RL (main text) or supervised learning (Appendix A.2.2) on several different tasks that are each defined by an underlying value structure: bandit tasks, reversal tasks, and hierarchical tasks (Figure 1e). For example, the reversal *task* has an underlying value structure in which the values of the two available

actions are anti-correlated (when one action is bad, the other is good), with these contingencies reversing at random. Each reversal *problem* is governed by this same structure but with *different available actions*.

We propose that irrespective of the structure of the value-based task, RNNs learn the same general circuit implementation to solve new problems within each task. This implementation consists of two components: 1) working memory (WM) subspaces and 2) value control signals (Figure 1d). The activity in the WM subspaces encodes the identities of the actions that are available in the current problem, with one action stored in each subspace. These subspaces are organised in order of preference, so that action representations are stored in ranked order (Figure 1d R1, R2, R3). The output action is the action stored in the ‘best’ (R1) subspace. To change behaviour in response to reward feedback and select a different output action, action representations can be dynamically re-organised between the WM subspaces such that a different action occupies the R1 readout subspace. Determining the precise re-ranking of actions between ranked WM subspaces requires tracking the values of each ranked preference (Figure 1d V1, V2, V3). Comparisons between these value signals gate the movement of action representations between WM subspaces. Critically, the value representations are *action invariant*, and instead encode the value of ‘best’ (V1), ‘second best’ (V2), and so on.

As an example, consider a 2-arm bandit problem where the possible actions are ‘watermelon’ and ‘banana’ (Figure 2b). Initially, the watermelon might be preferred over the banana, so the rank 1 subspace contains the watermelon action representation and rank 2 the banana representation. However, upon choosing the preferred action ‘watermelon’ and receiving little reward, this ranking may need to change. Changing action preferences happens by re-ordering the activities stored in each ranked subspace; ‘watermelon’ moves down to rank 2, and ‘banana’ moves up to rank 1 (Figure 2d). Determining when to re-order options requires tracking and comparing the values of each rank (Figure 2d). In the 2-arm bandit, switches occur only when the value of the current preferred action (rank 1) falls below that of the non-preferred action (Figure 2b), as illustrated in steps 4 and 5 of Figure 2d.

**Figure 2:**
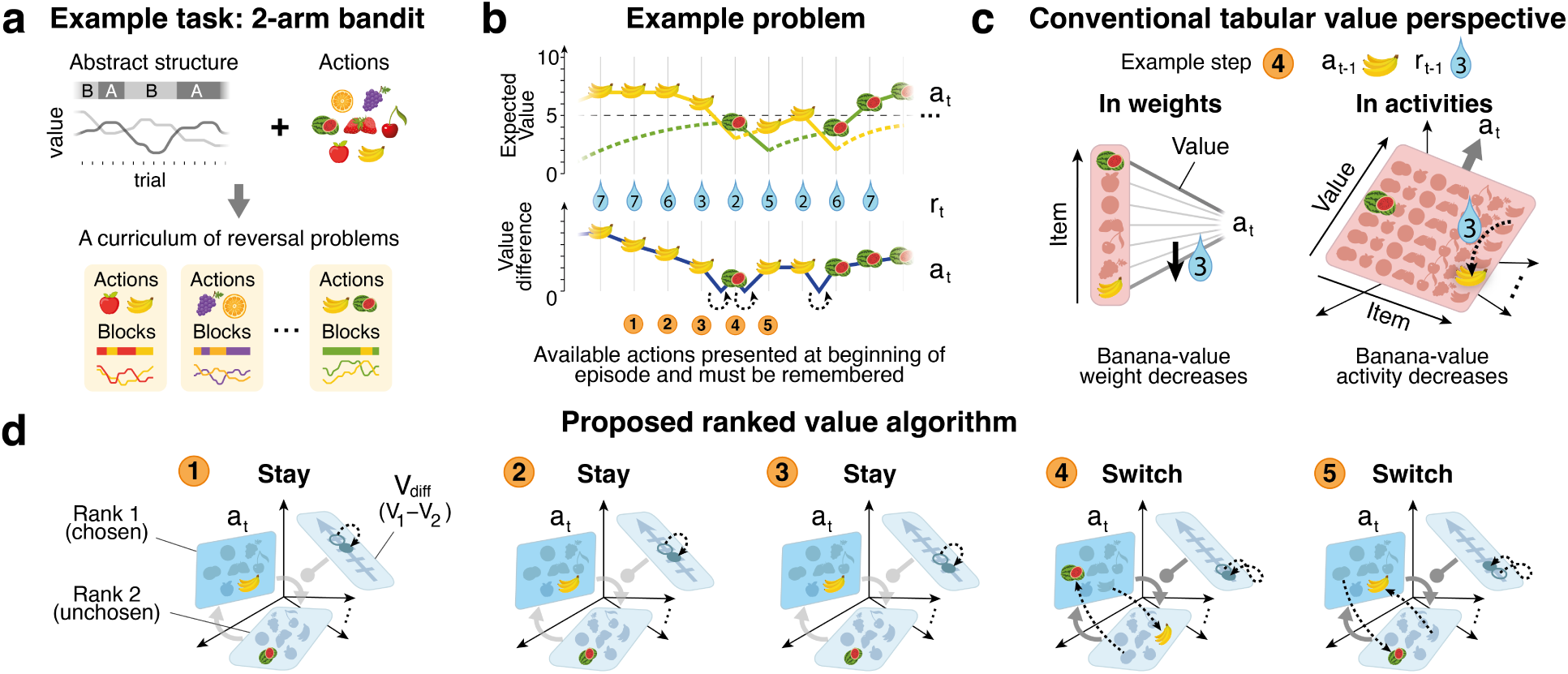
Proposed ranked value implementation on a 2-arm bandit task. **a)** We consider meta-RL tasks in which each problem is drawn from a task distribution composed from a consistent task structure and a set of actions. For illustration we show a generalised 2-arm bandit task where the abstract structure consists of two randomly drifting reward magnitudes. **b)** Example problem showing values of two options. Value of the chosen option exactly tracks reward magnitude received (*r_t_*_−1_), whereas unchosen value decays towards the mean. Determining when to switch choice requires computing the value difference (bottom). **c)** Conventional (tabular Q-learning) RL specifies that action-specific values are updated with reward feedback (here reward 3 after choosing banana). This could be implemented either in weights (left) or activities (right). **d)**Flow of information between subspaces for five consecutive trials from **b**. General (action-invariant) value axis tracks difference in value between ranks 1 and 2 (upstream representations for *V*_1_ and *V*_2_ have not been illustrated for clarity). Before step 4, no switch occurs as value difference is positive. In step 4 a switch occurs, since value of preferred option (*V*_1_) falls below value of un-preferred (*V*_2_) after receiving *r_t_*_−1_ = 3. Value difference rebounds to be positive, since the new choice is now more valuable. Contents of chosen and unchosen subspaces are swapped.

This mechanism allows value-integration rules to be re-used in new problems, since the value representations are separate from the action representations. To solve a new unseen problem, all that is required is for the actions relevant to a problem to be loaded into the working memory subspaces.

The proposed generalised value representations differ from the canonical descriptions of RL in the brain, which have been a successful models of striatal learning (Figure 2c). In the striatal description, values are directly tied to actions or stimuli via synaptic plasticity between cells encoding the action/stimulus, and cells encoding their value. Hence, value coding is tied to each specific stimulus or action, not the representations of a general preferred and second-preferred (etc.) action. This is limiting in two regards, first it means the same value update rule must be re-learnt every time the stimulus or action set changes, limiting generalisation of the learnt rule to novel actions. Second it means the learning algorithm itself is determined by evolution – in this case corticostriatal plasticity. As we will see, generalised value representations allow more efficient structured learning algorithms to be learned and embedded in the synaptic structure of the prefrontal cortex.

## 3 Results

### 3.1 Agent set-up

#### Task

We design several meta-RL tasks which test whether networks learn task structure and apply it to new problems (Figure 1e, Appendix A.1 for detail on each task). Each task consists of many sub-problems which share the same task structure, but have different actions available. The available actions in a problem are presented sequentially at the beginning of the episode. Each problem is itself a sequential decision making problem, in which an action must be chosen on each trial (timestep) and reward feedback is given after each action. To induce task structure, reward probabilities and magnitudes, for each action, vary through time according to a generative model that embeds specific task rules (see Figure 2a for the 2-armed bandit example). In order to maximise reward on held-out problems, the common structure underlying all problems must be learned on training problems, and generalised to new problems.

#### Network details

For our agent RNN, we use gated recurrent units (GRUs), with linear readouts from the actor (policy) and critic (value), and linear input embeddings. On each timestep, the previous action *a_t_*_−1_ and previous reward *r_t_*_−1_ are provided and the network must select an action *a_t_* (Figure 3a top). To model real-world choices where many actions are available, we allowed the agents to choose all possible actions, and simply receives no reward if that action is not ‘active’ in this problem.

**Figure 3:**
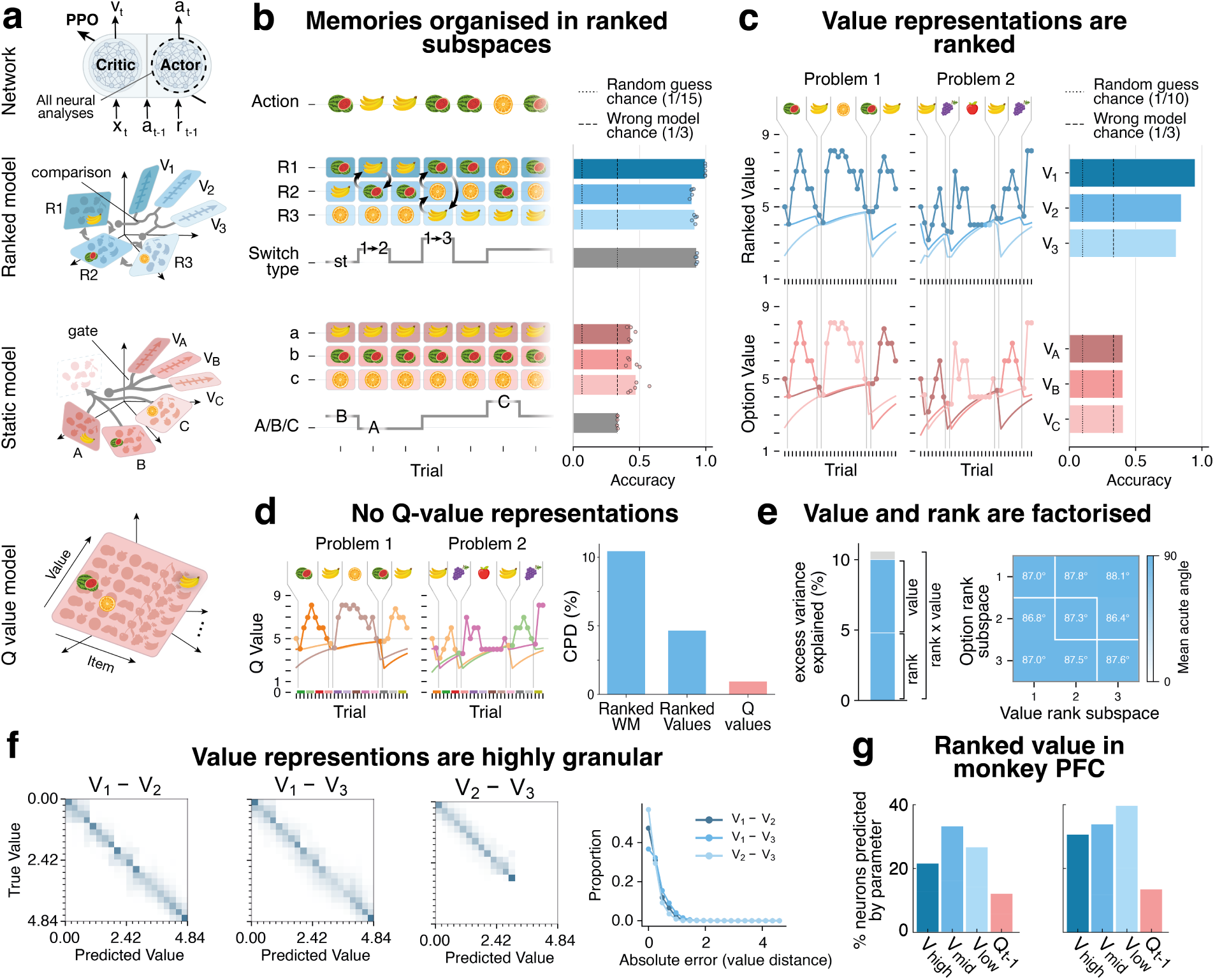
Network learns ranked WM solution on 3-arm bandit task. **a)** Top: network architecture - separate RNNs for actor and critic; all analyses performed on actor RNN. Below: three hypothetical learned models: proposed ranked WM model (blue, top), a model with static working memory and a gateable readout (red, middle), a Q-value model (red, bottom). **b)** The hypothetical contents of subspaces on an example problem. Blue shows ranked model, with options dynamically re-ranking between subspaces (blue cells), and corresponding control switch signals (blue trace). Red shows equivalent gated model. Contents of each of the three WM subspaces (from the set of 15 total actions) and switch signals, are decodable from actor RNN (blue bars) but not from gated model (red bars). **c)** Two example problems with putative ranked value traces (blue) and static value traces (red). The three ranked values are decodable from actor RNN using logistic regression (blue bars), while the static values are not (red bars). **d** Q-value traces for two example problems; unlike **c**, different representations are active in each problem (aside from the banana action value which is common to both problems). Q value has low CPD relative to the ranked signals. **e)** Left: ranked options and ranked value together explain nearly the same amount of hidden activity variance as the conjunctive ranked option x ranked value regressor, indicating the representations are factorised rather than conjunctive. Right: subspaces (obtained from decoders) for ranked options and ranked values are all approximately orthogonal. **f)** Confusion matrices show that differences between ranked values (required for behavioural switches) are decodable with high precision using logistic regression. **g)** Ranked value representations in Macaque PFC for two subjects, figure adapted from Knudsen & Wallis (2020). Panel **b** shows 5 separate training runs with different seeds, **c-f** show 1 run.

#### Training

To demonstrate that any learnt circuit mechanisms are not a consequence of training decisions, we use two different training regimes; RL (main text) or supervised learning (appendix). For the RL regime, to be certain that any value representations found in the actor are not a consequence of training the network to predict value, we therefore use independent networks for the actor (policy) and the critic (value). We analyse only the actor network. In RL, we train with proximal policy optimization (PPO; Schulman et al., 2017). For the supervised regime, we use the actions of an handcrafted optimal agent as targets for gradient descent. We train networks to convergence and test on a held-out set of unseen problems to ensure generalisation (Appendix A.2.3).

### 3.2 Interrogating candidate circuit mechanisms

While our proposed model (previous section) is sufficient to solve these tasks, there are other potential implementations (that RNNs or PFC could in theory learn) of how action memories are organised and how value determines action selection. We consider two alternative candidate implementations — static gated memory and Q-value. The three candidates are outlined below (Figure 3a).

First, in the **ranked working memory** solution (our proposed model), the network has memory subspaces that store actions in their preference order. Read-out is simply choosing the action in the preferred subspace. Value representations in this model are also ranked. For example, there is a subspace for the value of the best option and simultaneous (orthogonal) subspaces for the value of the second best option and so on. There is no representation of the value of a specific action (which is what conventional RL requires). In the ranked working memory model, value’s role is to *control the movement of action representations* between the action memory subspaces. For example, when the best value becomes lower than the second best value, the rank 1 and rank 2 action memories should be swapped.

Second, in the **static gated memory** solution there are also memory subspaces, but the content of the memory subspaces do not change through time. Each ‘active’ action in a problem is assigned to subspace (e.g., subspace A, B, or C). Value representations are tied to each memory subspace (”The value of action A”), such that the values can *gate which memory subspace is read out into the action subspace*. We define A, B, C as the order in which the available actions are presented at the beginning of the episode.

Lastly, we consider a conventional **Q-value** solution that tracks the value of each action. Here, there are no working memory subspaces, but instead there are value representations for each of the actions in the entire action set, which are directly modified as values change across trials. The chosen action is determined by the relative values across the action set. Hence, this solution has no generalised (action-invariant) value subspace, but instead value is specific to each actual action (Figure 3d).

To distinguish between these implementations in our trained RNNs, we seek to understand three things. First, if the network has a working memory representation of a problem’s available actions, and, if so, whether it is ranked (best to worst action) or static (actions A, B, C). Second, if the network (the actor RNN not the critic) represents value and whether values are tied to a rank (ranked working memory model), an available action A, B, C (static gated memory model), or a specific action (Q-value model). Third, how value subspaces causally influence the memory and output subspaces, as this differs in each implementation.

To answer these questions we first trained linear decoders on the (trained) actor RNN hidden activity to predict memory and value content in different frames of reference (Sections 3.3 & 3.4), and then performed causal interventions on the specific subspaces in the actor RNN (Sections 3.5 & 3.6).

### 3.3 RNNs learn a ranked working memory implementation on the bandit task

The first task structure we analyse is the 3-arm bandit task. Here, the task structure shared between problems is that there are always three action values that drift independently over time. In this task, the rewards change in magnitude and are non-probabilistically delivered. What’s different in each problem are the ‘active’ actions and the (random) drift. In both training regimes, the agents generalised to held-out problems with near optimal performance (Appendix A.2.3). We were therefore able to interrogate the trained models to understand the circuit implementation. All analyses were performed on the problem set that was held-out from network training (i.e. novel triplets of available actions).

#### 3.3.1 Working memory subspaces encode ranked actions

We first tested the organisation of the actions in working memory and found that this favoured the ranked working memory implementation over the static gated memory implementation. We generated hypothetical contents of the three working memory subspaces in each model. For the ranked model R1 contents were defined as the current action taken, R2 as the next action following a swap, and R3 as the remaining action (Figure 3b blue cells containing ‘fruit’ memories); for the static model, A, B, and C contents were defined as the actions in the order they were presented, and kept static throughout the duration of the episode (Figure 3b red cells). We could decode the hypothesised actions stored in each of the three ranked WM subspaces (Figure 3b blue bars: R1, R2, R3) but could not decode the hypothetical contents of the static gated memory implementation (Figure 3b red bars: a, b, c). Chance levels were 1*/*15 for random guessing between all available actions, and 1*/*3 for wrong memory frame-of-reference.

We then tested how memories were routed to output the action. For the ranked working memory model, the readout projection is static while the action memories are dynamically re-organised between the subspaces to place the correct action in the R1 subspace. This requires a control signal encoding the type of swap (grey round-headed lines in Figure 3a ranked model). The three types are best moves to middle (1 → 2), best moves to worst (1 → 3), and best stays best (Figure 3b grey ‘switch type’ trace). For the static working memory model, the action memories are static while the readout is dynamically shifted between the subspaces, routing the memory to output. This requires a control signal encoding which subspace to readout from (Figure 3a static model, Figure 3b grey ‘A/B/C’ trace). We could decode the switch control signal in the ranked working memory model but not the static gated model (Figure 3b).

#### 3.3.2 Value representations in ranked coordinates

Next, we tested how value was encoded. Because reward is only delivered for the chosen action (i.e., there is no feedback for 2 of the 3 drifting values), we estimated the value of the two non-chosen actions by decaying their value towards the mean midpoint value, from the trial that it was last sampled (see methods A.1.4 for theoretical motivation and details). We were able to decode the binned value (10 bins) of each of the three ranks (value of best, middle, worst) but not of each of the three available actions (value of A, B, C) (Figure 3c), supporting the ranked over the static working memory model. Balanced class weights were used during decoder training, giving a chance level of 1*/*10 for random guessing between bins and 1*/*3 for the wrong value frame-of-reference. To interrogate whether Q-value representations were present, we used a variance-based analysis rather than decoding analysis. This is because if a particular action is high value, that action is more likely to appear in the top rank of a ranked memory model and so a decoding analysis is not sufficient to distinguish between the two models. Instead, we calculate the unique variance in hidden activity explained by the ranked values and Q values (Figure 3d right), after controlling for nuisance variables (see methods). This favoured ranked value representations over Q-value. We note that ranked value representations mirror neural recordings from macaque PFC (Knudsen & Wallis, 2020) in which a larger proportion of neurons code for value of best, middle and worst options (*V_high_, V_mid_, V_low_*) than for item value (*Q_t_*_−1_) (Figure 3g).

To further confirm that ranked value is generalised, i.e. factorised from ranked action, we used two methods. First, we calculated the variance of hidden activity explained by ranked working memory contents and ranked values as separate regressors, in comparison to their conjunction (interaction) as a single regressor. We found that the conjunction of ranked value and ranked action resulted in little gain in variance explained compared to the two factors separately (Figure 3e left). This implies a factorisation of ranked working memory and ranked value. Second, we measured the angles between the decoder readout weights for the three ranked working memory representations and ranked value representations, finding that they were all approximately orthogonal (Figure 3e).

Finally, we interrogated the nature of the ranked value representations and found that they were highly granular, akin to a discrete code (for example, a place code) rather than scalar code (for example, a ramping signal). Binned value (Figure 3c) and binned value differences (Figure 3f) were decodable with logistic regression, implying a high-dimensional value code that partitions neighbouring value ranges into distinct representations. For value difference decoding we found a strong diagonal on the confusion matrices and relatively small absolute error (Figure 3f right). Notably, the difference between *V*_2_ and *V*_3_ was highly decodable, despite the two values being highly correlated. Discretely partitioning the value axis into distinct representations in this manner is important, as it allows for neighbouring values on either side of a decision boundary to exert different effects on downstream computations.

### 3.4 RNNs learn ranked working memory implementations on structured value tasks

The bandit task involves choosing between actions that have uncorrelated independently evolving values, however this is often not the case in real-world decisions. It is common for the reward contingencies of different actions to relate to each other in a structured manner, such that knowing the value of one action is informative of the other values (Figure 4a, c). If these relations between values are understood, then the best action can be inferred without ever needing to sample it. This allows for much more sample-efficient behaviour.

**Figure 4:**
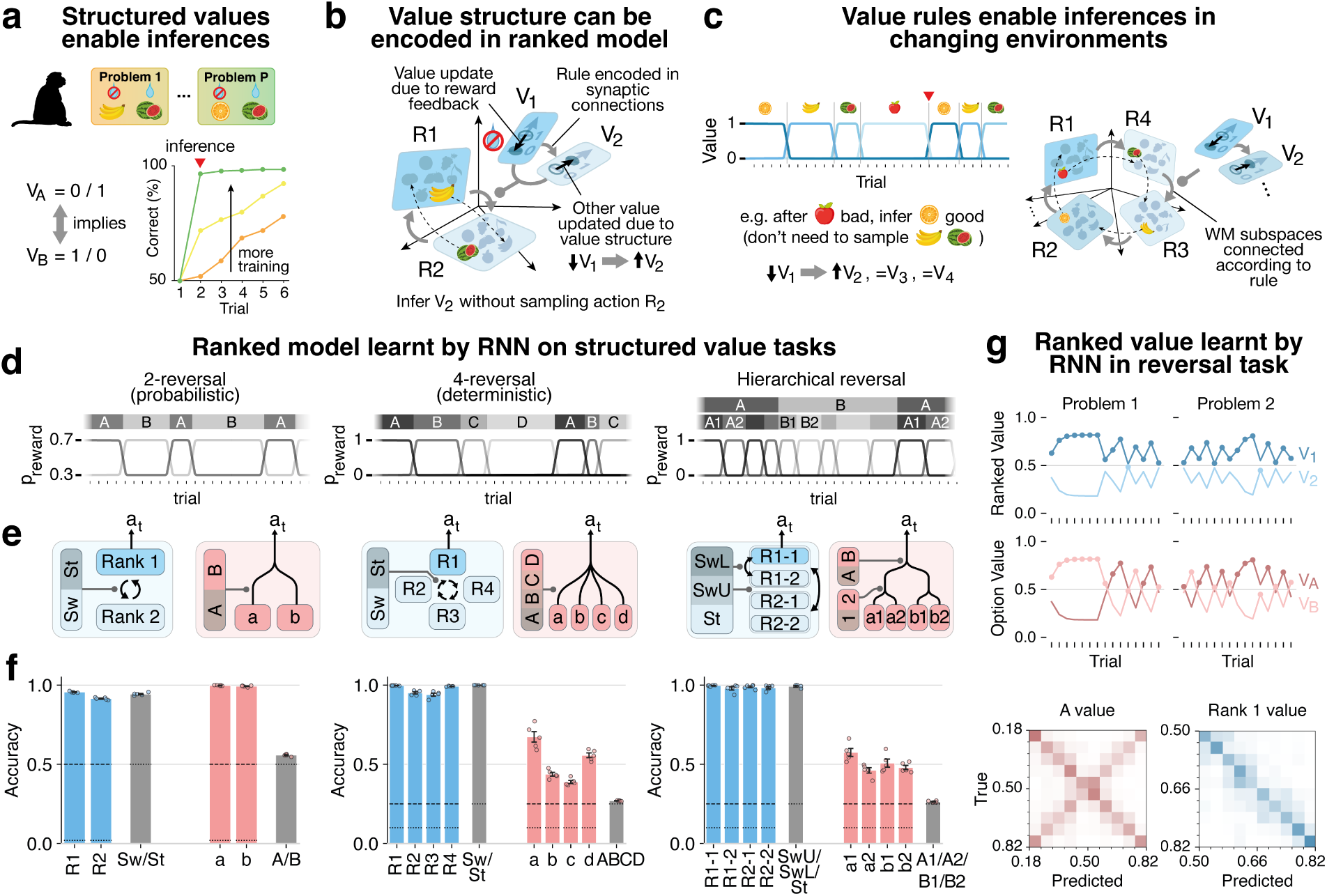
Ranked WM model can encode structural relations between values. **a)** An example task (Harlow, 1949) with a value rule - one item rewarded and other unrewarded. In each problem animals are shown two items, and can choose freely between them over several trials. After training (green trace), one only needs to sample one action to infer the value of the other (and therefore the correct action). Children, monkeys (Harlow, 1949), and rats (Slotnick & Katz, 1974) exhibit one-shot inferences on the task. **b)** The corresponding value rule (here, the Harlow rule i.e., anti-correlated values) is easily encoded in a ranked model. After choosing banana in problem 1 and being unrewarded, the chosen value (*V*_1_) is updated to 0. By exploiting the structure, *V*_2_ can simultaneously be updated to 1, even without sampling the corresponding action. This is mediated by synaptic connections between the value subspaces, which embed the rule. A switch occurs as *V*_2_ *> V*_1_, and the animal chooses watermelon on the next trial. Note that in the case of deterministic reward, no working memory value representations are required and reward can be used directly as the switch signal; value was illustrated for narrative clarity. **c)** Value structure can be temporal, for example with values evolving in a cyclic order (top). This enables the value of unsampled options to be inferred and rapid adaptation when contingencies change. This value rule can be embedded in the ranked working memory solution via the connections between subspaces. **d)** Three structured value tasks, in which values evolve co-dependently. **e)** Contents of ranked WM (blue) and switch signals (grey) are decodable from the actor RNN of networks trained on each of the three tasks, whereas signals from the static models are not (red) (protocol as in Figure 3b). **f)** In the probabilistic reversal task networks must integrate reward to track value or belief. Chosen (rank 1) value (blue traces) obtained from an ideal Bayes agent is decodable with high granularity from the actor RNN whereas state A/B value (red traces) is not. Note that chosen value is analogous to belief that the current action is best. Panel **e** shows 5 separate training runs with different seeds, **f** shows 1 run.

The ranked working memory model enables these action inferences. This is because the structural relations between values can be learnt in the synaptic connections between the value representations, allowing for unsampled values to be inferred and therefore for unsampled actions to be re-ranked in working memory according to the updated preferences. For example in a Harlow-like task in which there are two anti-correlated action values (Figure 4a), after taking an action and finding out it is low value, one can deduce that the other action is high value. In a circuit, this could be implemented via synaptic connections between the representation encoding low *V*_1_ and the representation encoding high *V*_2_, such that activity in one induces activity in the other^1^. In this way, value feedback for the chosen action (*V*_1_) can be used to update the value of the unchosen action (*V*_2_). These inferred values can then be used to control the switching of WM contents (Figure 4b). Since the value representations are factorised from the action representations, the value rule (embedded in the connections between value representations) can be applied to new contexts that share the same rule but have different available actions (Figure 4c).

To test whether the ranked working memory implementation is learnt on structured value tasks, we trained (separate) networks on three different tasks (Figure 4e top row), using either reinforcement learning (main text) or supervised learning (Appendix A.2.2). Each task has a value rule which is shared across problems, with different available actions on each problem.

#### Task 1: probabilistic reversal task

(Figure 4d left) in which there are two actions with anti-correlated reward contingencies (*p_reward_* = 0.7 for current best action, *p_reward_* = 0.3 for worse action). These contingencies swap synchronously and at random in a block structure. Reward feedback is probabilistic, so rewards need to be integrated over time in order to compute a value or belief of the best option.

The ranked working memory implementation for the reversal task has two action memory subspaces— chosen (R1) and unchosen (R2) (Figure 4e left blue model). Rank 1 (chosen) value is tracked (Figure 4f dark blue traces) and controls when to reverse the actions stored in the two subspaces; when chosen value is sufficiently low and no reward is received, a swap must occur. There is no need to track rank 2 value as it is exactly anti-correlated with rank 1 value. The static gated memory implementation has two subspaces, A and B, storing the two available actions in the problem, with readout gated by a control signal encoding which of A or B is better (Figure 4e left red model). Value of A is tracked to compute this control signal (*V_B_* not needed as it is exactly anti-correlated).

We were able to decode the contents of the two ranked working memory subspaces, as well as the reversal signal for swapping the contents (Figure 4e blue bars and grey bar respectively). While we were also able to decode the contents of A and B subspaces, we could not decode the A/B control signal that would be required to use these to control action output (Figure 4e red bars and grey bar respectively).

We were also able to decode ranked (chosen) value but not A value, consistent with the ranked over static model. When decoding *V_A_* we found an ‘X’ shaped confusion matrix, implying that the decoder confused high *V_A_* with high *V_B_* (low *V_A_*), and therefore that the representations for each are similar (Figure 4f). This implies a chosen value frame of reference.

#### Task 2: cyclic reversal task

(Figure 4d middle) in which there are 4 values which each become ‘good’ in a predictable sequential order, but at random intervals. When the current best action becomes bad (*p_reward_* = 0), the next action in the loop becomes good (*p_reward_* = 1). Note that reward is deterministic so the agent does not need to integrate value over time to infer the best option.

A ranked working memory implementation for the cyclic task would consist of four ranked action memory subspaces *connected in a loop* (*R*4 *R*3 *R*2 *R*1 *R*4 …) and a signal controlling whether to ‘turn’ the four memories around the four subspaces (Figure 4e middle blue model). These connections between the ranked subspaces embed the rule. A static implementation consists of four action memory subspaces (A, B, C, D) and a gating signal (A/B/C/D) (Figure 4e middle red model).

We could decode the contents of the four ranked working memory subspaces but not of the A, B, C, D static subspaces (Figure 4e middle blue and red bars), consistent with the ranked model. While decoding the switch signal for the ranked model is trivial (reward input is the swap signal), we could not decode an A/B/C/D control signal (Figure 4e middle grey bars).

#### Task 3: hierarchical reversal task

(Figure 4d right) in which there are 4 values grouped hierarchically into two sets of two. At any time, one set is ‘best’ and the two values within this set alternate at random intervals, as in a reversal task (*p_reward_* = 1 for the best action, *p_reward_* = 0 for the worse within-set action, *p_reward_* = 0 for the other actions). Which set is best alternates on a longer timescale at random intervals. This requires hierarchical inference of which set is currently best, and which action within the set is best. Reward is deterministic, so the network does not need to integrate value over time, but it does need to count consecutive absences of reward to infer the best option (1 miss is a within-set swap, 2 misses is a set swap).

The ranked working memory implementation (Figure 4e right blue model) has four ranked action memory subspaces organised hierarchically into an upper set (R1-1, R1-2) and a lower set (R2-1, R2-2), with a control signal selecting between two possible swaps. The two possible swap types are a lower-level swap in which only the actions in the current set are swapped (R1-1 R1-2), and an upper-level swap in which the two whole action sets are swapped ((R1-1, R1-2) (R2-1, R2-2)). This requires within-set connections gated by 1 reward miss, and between-set connections gated by 2 reward misses. We tested for a hierarchical static implementation, with four working memory subspaces (A1, A2, B1, B2), and a gating signal for selecting which of A1/A2/B1/B2 to select.

We could decode the contents of the four hierarchically ranked working memory subspaces but not of the A, B, C, D static subspaces (Figure 4e right blue and red bars), consistent with the ranked model. It was also possible to decode the type of switch (‘stay’, ‘switch lower’, or ‘switch upper’).

In summary, we observed that networks trained on structured value tasks recovered ranked working memory implementations analogous to that found for the 3-arm bandit task. We observed this both in the reinforcement learning regime (main text) and the supervised learning regime (Appendix A.2.2). Furthermore, ranked value was found in networks trained on the reversal learning task, which required integration of reward into value (or belief). This parallels the representations found in the 3-arm bandit task, despite the difference in the nature of the value feedback — in the bandit task varying reward magnitudes were deterministically provided by the environment, while in the reversal task binary rewards were probabilistically provided.

### 3.5 Ranked WM subspaces causally determine future behaviour

#### 3.5.1 Causal intervention using linear encoders

To prove that the identified ranked working memory subspaces and ranked values are functionally relevant for behaviour, we performed causal interventions on the hidden activity of the actor RNN. In particular, we performed targeted manipulations of these variables in the hidden activity and observed the effect on both future internal representations and future behaviour. To achieve this, we trained linear encoders to map from a candidate variable (for example, the hypothesised rank 2 working memory content, Figure 5a) to the hidden activity of the actor RNN (Figure 5b left). Once trained, this encoder allowed us to remove activity relating to the true state of this candidate variable, and replace it with a manipulated state (Figure 5b right, activity relating to 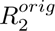 removed and replaced with 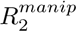). We used this method to manipulate working memory (Section 3.5) and value (Section 3.6)

**Figure 5:**
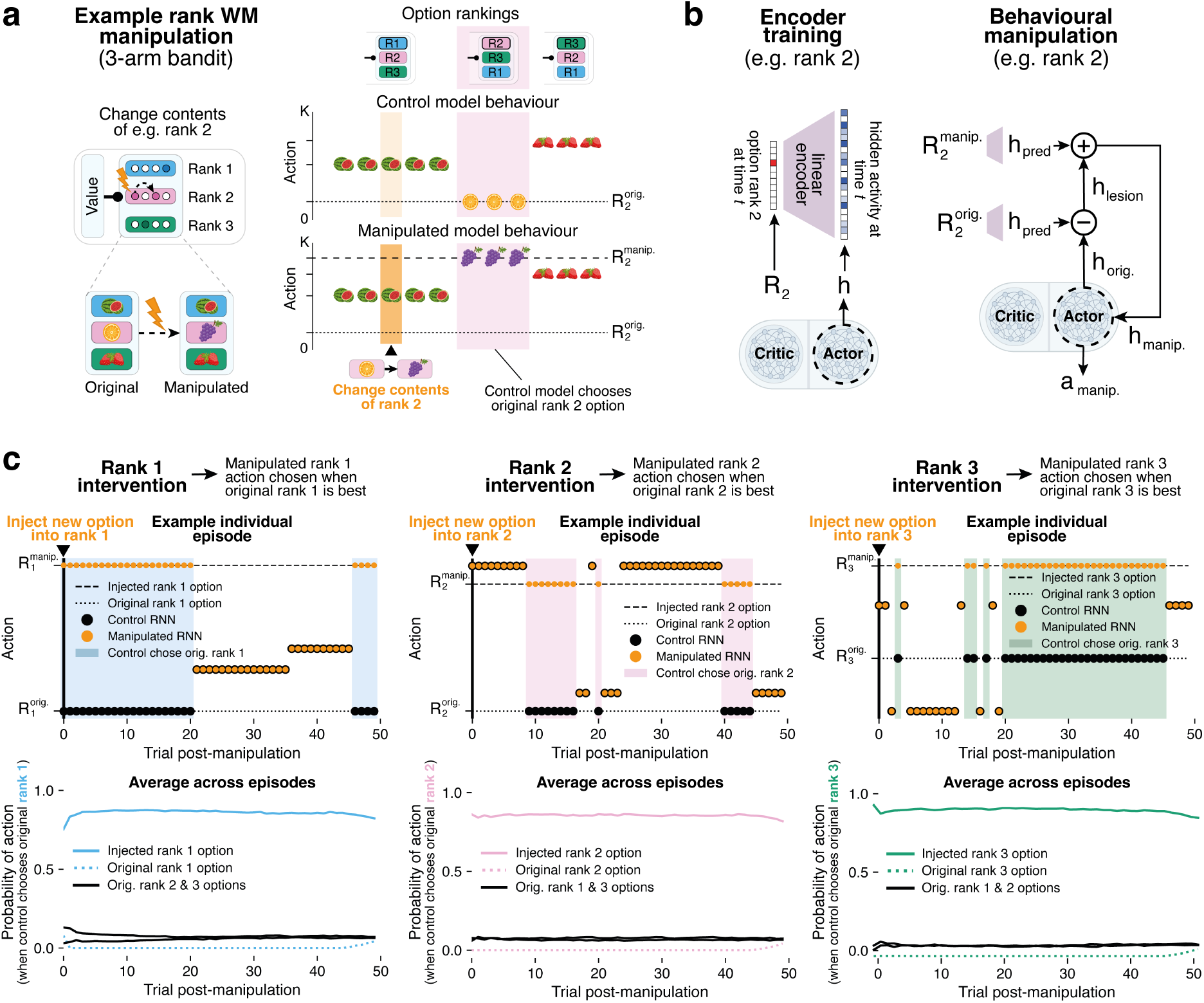
Causal manipulations of ranked action subspaces predictably affects future behaviour. **a)** Hypothetical manipulation of contents of rank 2 WM subspace. Left: representation of rank 2 option (orange) is swapped for another option (grape). Right: hypothetical behaviour of control (un-manipulated) model and model with manipulated rank 2 memory. Manipulated model behaviour mirrors that of control model, but chooses injected rank 2 option (grape) whenever control chooses original rank 2 option (orange). Top shows time course of WM rankings (relative to order at time of manipulation). **b)** Protocol for injecting new option representations into WM subspaces of the actor RNN (rank 2 example shown). Left: a linear encoder was trained to predict actor hidden activity from option currently rank 2. Right: replacement of original rank 2 option representation performed via lesion (removing hidden activity associated with current rank 2 option 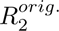) followed by injection (adding hidden activity associated with new rank 2 option 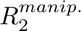). Network is then simulated with new manipulated hidden activity. **c)** Top: behaviour in example episode following manipulation reproduces that of panel **a**. For each rank manipulation, when target rank is best (shaded pastel regions) the manipulated RNN chooses the injected option rather than original option corresponding to that rank. Bottom: in shaded regions only (original target rank becomes best) across all episodes, probability of manipulated RNN choosing injected option is high.

#### 3.5.2 Causal WM intervention procedure

Causal manipulation was performed for each of the three ranked working memory subspaces in the 3-arm bandit task, using three separate encoders, one trained for each ranked WM subspace. At a random timestep, the content of a ranked working memory subspace was changed. According to the ranked working memory implementation, the effect of this manipulation on subsequent behaviour should be that the newly injected action is chosen whenever the control (non-manipulated) model would have chosen the action originally stored in the targeted rank. There should be no effect on the contents of other memory subspaces and therefore other actions should be identical for both models. For example, if the contents or rank 2 are swapped from orange to blackberry, then whenever the control RNN chooses orange, the manipulated RNN should choose blackberry (Figure 5a right). When the control network chooses the actions that were rank 1 and rank 3 at time of manipulation (watermelon or strawberry), the manipulated model should choose the same actions. Note that when injecting a new action representation into the network, we simultaneously updated the environmental available action set (i.e., the problem) so that the environment started delivering rewards for the injected action and stopped delivering rewards for the erased action.

#### 3.5.3 Injecting new action representations into ranked WM subspaces induces selection of new actions

We found that the behavioural effect of manipulating ranked working memory contents was as predicted. When manipulating the rank 2 subspace (scenario illustrated in Figure 5a), this does not affect the immediate behaviour because the network chooses the rank 1 action. However, later on when the control network chooses the action originally in rank 2 (pink shaded region, Figure 5c top-middle), the manipulated network instead chooses the action that was injected into rank 2 (orange dots). This means that the action memory that was injected into the rank 2 subspace was later moved into the rank 1 subspace. When the original rank 1 and 3 options are best, they are chosen by both the control network and the manipulated network since only the original rank 2 subspace contents were manipulated. Similarly for rank 1 (Figure 5c top-left) and rank 3 (Figure 5c top-right) manipulations, the manipulated actor RNN subsequently chooses the manipulated action only when the control actor RNN chooses the original rank 1/3 action (blue/green shaded region), consistent with predictions from our ranked WM subspace model.

The behavioural effect post-manipulation was long-lasting, meaning that the pre-manipulation memories were fully eradicated and replaced. To summarise this, we took the trials in which the control RNN chose the action originally contained in the rank being targeted for manipulation (blue/pink/green shaded regions) and computed the fraction of times that the manipulated RNN instead chose the injected action (Figure 5c bottom plots). We observe that, post manipulation, there is a high likelihood of the injected action being chosen instead of the original action, and this effect lasted indefinitely throughout the episode.

We performed an equivalent manipulation of ranked working memory contents in a network trained on the probabilistic reversal task and observed an identical effect (Figure 10), demonstrating that ranked memory slots causally determine behaviour in different value tasks. Since static memory slots (A and B) were also decodable in the reversal task, we performed an intervention in which the contents of A and B subspaces (Figure 4e left red bars) were swapped. If these were used in a static gated memory implementation to causally determine action output, then swapping the contents would swap the two actions in behavioural output. Post-intervention behaviour was unchanged (Figure 10c), meaning A and B subspaces were not causally linked to which of the two actions is selected. This is in agreement with the absence of an A/B gating signal (Figure 4e), which is also required for a static gated memory implementation.

### 3.6 Ranked value representations control ranked working memory contents and behaviour

#### 3.6.1 Causal value intervention procedure

We performed equivalent manipulations on the ranked value representations to determine whether they (1) dynamically control the re-organisation of action memories in the ranked WM subspaces and (2) subsequently determine the actions selected by the network. To manipulate activity, we followed an analogous protocol to that described in Section 3.5.1 by first training a linear encoder to map (binned) ranked value to hidden activations of the actor RNN. This allowed us to change the value state of the actor RNN by first erasing and then injecting value-related activity.

In the 3-arm bandit task, there are two critical value thresholds, corresponding to when rank 1 value becomes smaller than the two other ranked values: if *V*_1_ falls below *V*_2_ (*V*_2_ − *V*_1_ *>* 0), and if *V*_1_ falls below *V*_3_ (*V*_3_ *V*_1_ *>* 0). According to the ranked working memory implementation, the former should lead to the action memory in rank 1 moving to rank 2 (illustrated in Figure 6b), and the latter leading to a complete reconfiguration of the memories: rank 1 action memory moving to rank 3, rank 3 moving to rank 2, and rank 2 to rank 1.

**Figure 6:**
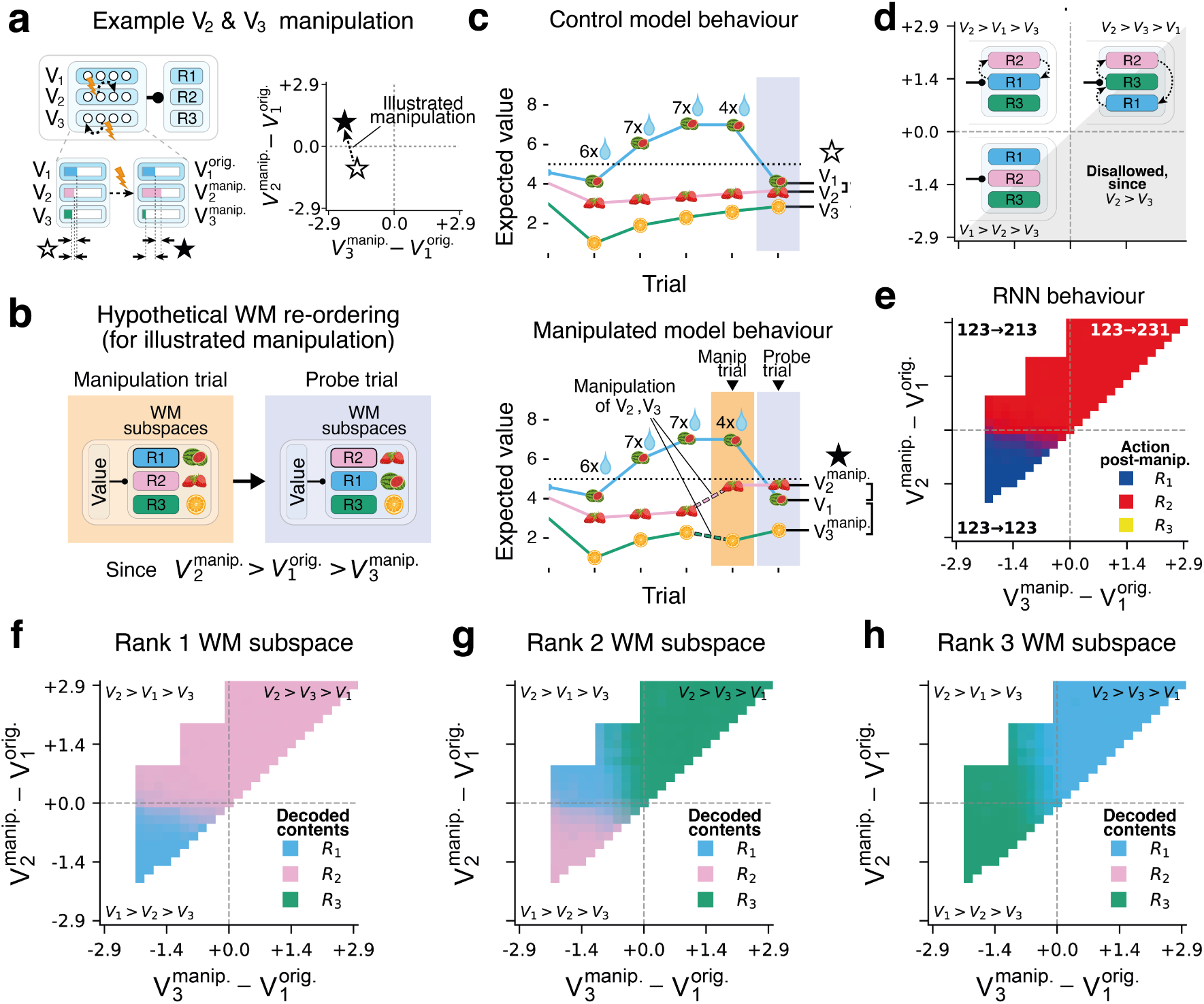
Causal manipulations of value in actor RNN re-ranks action subspaces and predictably affects future behaviour. Manipulation of value-related activity in actor RNN for probabilistic reversal task and 3-arm magnitude bandit task. **a)** Left: schematic of manipulation targeting value subspaces. Curved dotted arrow between neurons in the model denotes change from original value to new manipulated value. Right: Solid star and empty star denote pre- and post-intervention value respectively for the illustrated manipulation on the left, plotted within the 2D space of all possible manipulations. **b)** Illustrated value manipulation triggers hypothetical re-ordering of contents of WM subspaces on subsequent trial. R1/2/3 (blue/pink/green) denote the option stored in rank 1/2/3 WM subspaces just prior to manipulation. **c)** Top: hypothetical behaviour of control (un-manipulated) model - in illustrated cases no switches are observed. Bottom: hypothetical behaviour of manipulated model - R2 (strawberry) becomes more valuable than R1 (watermelon) so is selected post-intervention. **d)** The 3 quadrants of behaviour (bottom left *V*_1_ *> V*_2_ *> V*_3_ i.e. 123 123, bottom left *V*_2_ *> V*_1_ *> V*_3_ i.e. 123 213, *V*_2_ *> V*_3_ *> V*_1_ i.e. 123 231). Grey shaded region denotes value space that is unreachable (*V*_2_ *> V*_3_, since *V*_2_ and *V*_3_ are never directly sampled and so cannot switch). **e)** Manipulation predictably affects RNN behaviour: when 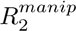 > *V*_1_, R2 (red) is chosen on the post-intervention trial, otherwise R1 (blue) is chosen. **f-h)** Memories are moved between ranked subspaces when manipulating ranked value across critical thresholds. In particular, see rank 2 WM subspace: each value quadrant results in different contents due to different swap types (bottom left, stay: *R*1*/***R2***/R*3, top left, best becomes middle: *R*2*/***R1***/R*3, top right, best becomes worst: *R*2*/***R3***/R*1, bold indicates rank 2 content).

We performed a sweep, where we replaced the original values with every other possible combination of values. In the 3-arm bandit, this required a two-dimensional sweep over every new combination of *V*_2_ and *V*_3_ (*V*_1_ could not be manipulated as it equals reward received as input). For each combination in the sweep we removed and injected activity corresponding to the representations of the new values 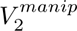 and 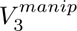. On the following trial, we observed whether (1) the action contents of the ranked working memory were re-organised in a manner congruent with the hypothesised induced switch type, and (2) the action taken by the network was congruent with the switch type. To determine the contents of the working memory subspaces, we used decoders trained to predict the ranked actions (as in Section 3.3).

#### 3.6.2 Injecting values around critical thresholds induces re-ranking of action representations in WM and behavioural switching

We found that the ranked value representations controlled both the output actions and the working memory action content in a manner exactly consistent with the ranked WM model hypothesis. When sweeping 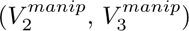 combinations, three quadrants emerged bounded by the two critical value thresholds:

##### Bottom left quadrant

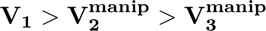. Ranked values remain in same order following manipulation, therefore no swap of working memory contents occurs — on the next trial the network selects the action that was rank 1 pre-manipulation (*R*1), the rank 1 subspace contains the action originally rank 1 (*R*1), rank 2 subspace contains the action originally rank 2 (*R*2), and rank 3 subspace contains the action originally rank 3 (*R*3) (Figure 6e-h bottom left quadrant).

##### Top left quadrant

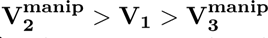. Ranked values are re-ordered, and we observe the corresponding re-ranking of action representations in working memory. Specifically, the contents of rank 1 and rank 2 WM subspaces are swapped — on the subsequent trial rank 1 WM subspace now contains action that was rank 2 pre-manipulation (*R*2) and rank 2 WM subspace contains the action that was rank 1 (*R*1), while rank 3 WM contents are unchanged (*R*3). The network outputs the pre-manipulation rank 2 action *R*2, indicating that a behavioural swap was induced (Figure 6e-h top left quadrant).

##### Top right quadrant

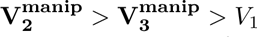. A complete re-ranking of actions occurs as best becomes worst and all actions move upwards one preference level. Rank 1 contains *R*2, rank 2 contains *R*3, rank 3 contains *R*1, and the network outputs the new best action *R*2 (Figure 6e-h top right quadrant).

The fourth quadrant (indeed the entire lower right triangle) does not exist, since in natural behaviour feedback is received only for the chosen action (*V*_1_). As the unsampled values (*V*_2_*, V*_3_) decay towards the mean, *V*_3_ can never overtake and be larger than *V*_2_.

We performed equivalent manipulations on the value representations of a network trained on the reversal task, and likewise found that value representations causally determined both the swapping of memories between the ranked working memory subspaces and the subsequent action output (Figure 11).

## 4 Discussion

We have presented a mechanistic model of how PFC solves flexible decision making problems, in which generalised value representations rank the available actions—via dynamic working memory representations— within a particular problem. We confirm this *in silico* across a wide range of tasks by showing that RNNs learn this exact model after training with either meta-reinforcement or supervised learning (Appendix A.2.2).

Critically, the learned values are unlike classical RL—not simply a Q-value output signal from which a policy can be read—but instead control signals that dictate the content and dynamics of working memory subspaces that contain action commands. The value signals proposed in our model for swapping memories between WM subspaces, and learned by RNNs, are consistent with PFC neural data: chosen (rank 1) value neurons (Padoa-Schioppa & Assad, 2006, 2008), unchosen (rank 2) value responses (Boorman et al., 2011), and best/middle/worst (rank 1/2/3) value neurons in Macaque PFC (Knudsen & Wallis, 2020). These cells generalise to different presented options (Padoa-Schioppa & Assad, 2008).

Our proposed ranked WM subspace structure for flexible decision-making closely resembles recent theoretical understanding of cognitive maps in PFC. In particular, in sequential working memory tasks (non-RL), PFC (Panichello & Buschman, 2021; El-Gaby et al., 2024) and RNNs (Whittington et al., 2025) structure memories of observations in relative co-ordinates (now, next, next next, etc., analogous to ranking) as opposed to absolute co-ordinates (A, B, C, etc.). In particular, one subspace encodes the ‘current’ memory of what is being observed, and all other subspaces encode the counterfactual memory of what would be observed at each relative location. An ‘action’ signal controls movement in the external world, and correspondingly controls the movement of memories between subspaces so that each subspace contains the correct relative memory. This analogy between how RNNs and PFC solve sequence memory (cognitive map) problems and flexible decision making problems, raises the tantalising possibility that structured working memory representations controlled by external or internally generated signals can offer an explanation to the mechanics of how PFCs solves increasingly general problems.

While we have framed much of our paper in terms of meta-RL, where PFC itself is trained to maximise reward, we also show that RNNs learn the same ranked working memory model when trained via supervised learning on optimal behavioural trajectories (Appendix A.2.2). Not only does this show that the ranked working memory model is a general solution to (meta-) decision making tasks, but it could also suggest an alternative learning paradigm for prefrontal cortex in which frontal cortex learns from optimal behavioural sequences replayed from an alternate brain region. For example optimal behaviours could be learned for individual problems by Q-learning in striatum, and these sequences could be replayed (via hippocampal memories) into prefrontal cortex which learns the general structure across problem sequences. The supervised regime is anatomically appealing, since reward feedback is used only as observation input to drive the network, in contrast to the meta-RL regime in which reward feedback must also train the weights themselves which requires dense dopamine projections to PFC—something that is not observed.

Our ranked working memory model contrasts with an influential idea for value-based decision-making: tracking the values of task latent states (Wilson et al., 2014). The latent state framework, e.g., in a 2-arm reversal task, suggests representing a variable for when A is good and similarly when B is good. Thus decisions become an inference over the underlying latent variable (i.e., which state you are in; A or B). When translated into the meta-learning setting, this requires additional bindings from latent state to action, since each new problem requires a different mapping from latent state to action. This could be done with synaptic memories (connections from state to action) which requires rapid learning, or with activity-based memories (subspaces A and B contain an action representations) as in the static gated memory implementation. It is interesting that RNNs did not learn this mechanism to solve our tasks—trained by either reinforcement learning or supervised learning—demonstrated by the inability to decode the current latent state i.e. whether the network is choosing A or choosing B, and the inability to decode static memory subspaces.

The value working memory model can be extended to offer plausible explanations for other puzzling properties of value signals in PFC. First, a striking property of PFC value representations is their strong within-trial dynamics. Individual value neurons have a preferred times within a trial to fire, so that the value population collectively tiles the entirety of the trial across pre-choice, post-choice and post-reward (Kennerley et al., 2009; Padoa-Schioppa & Assad, 2006; Samborska et al., 2021). Value representations gating the movement of working memory representations would provide a functional explanation for this. Since value must be maintained throughout the trial, and re-ranking of actions must occur only once per trial before choice time, it is necessary to have at least two value representations: one that is ‘potent’ and another that is ‘null’ with respect to gating working memory re-organisation. These null and potent subspaces would exchange activity in order to temporally orchestrate gating and subsequent behaviour, which parallels the potent and null subspaces in models of motor planning and execution (Kaufman et al., 2014; Elsayed et al., 2016). Second, in Neuroeconomic tasks in which subjects choose between two offers, during deliberation OFC has been observed to flicker between representing each of the two available options (Rich & Wallis, 2016). It has been proposed that this flickering is an integration or evaluation process, potentially sampling imagined or memorised values for each option (Balewski et al., 2023; Chen et al., 2026). In a ranked WM model this flickering would arise naturally, with chosen value integrating these noisy value samples over time, and rank 1 and 2 contents flipping during integration depending on which is currently better. We leave these directions to future work.

Prefrontal Cortex is critical to higher order cognition, with patients exhibiting profound deficits in behavioural flexibility should it be damaged (Funahashi et al., 1993; Bauer & Fuster, 1976). While PFC’s role in flexible decision making is well studied—both in behaviour and representation—there has been no mechanistic account that can unify the various roles and representations. By studying flexible behaviour more broadly, in particular meta-learning, we have shown that value and structured representations are two sides of the same coin, with value controlling the flow of structured representations, in line with the influential hypothesis that PFC is an executive controller of downstream representations (Miller & Cohen, 2001). While we have so far considered simple structured behaviours, we envisage that that our uncovered primitives, along with others, may be useful in extending our mechanistic understanding of flexible behaviour more broadly.

## Acknowledgements

We thank the following funding sources: Sainsbury Wellcome Centre’s core provided by Wellcome (219627/Z/19/Z; J.W) and the Gatsby Charitable Foundation (GAT3755; J.W., T.E.J.B); a Wellcome Principal Research Fellowship (219525/Z/19/Z; T.E.J.B); Gatsby Initiative for Brain Development and Psychiatry (T.E.J.B); the Jean Francois and Marie-Laure de Clermont Tonerre Foundation (T.E.J.B); Sainsbury Wellcome Centre’s core provided by Wellcome (219627/Z/19/Z; T.E.J.B); Sir Henry Wellcome Post-doctoral Fellowship (222817/Z/21/Z; J.C.R.W); European Research Council Starting Grant (NARFB/101222868; J.C.R.W); The Worshipful Company of Scientific Instrument Makers (J.W).

## A Appendix

### A.1 Task implementation

#### A.1.1 Problem structure

Networks were trained on one set of problems and evaluated on a held-out set. All permutations of each action combination (e.g. (*a, b*) and (*b, a*)) were assigned to the same split. The target train–test ratio was approximately 0.8:0.2, with approximately balanced coverage of individual action identities to reduce action-specific biases. Figure 9a illustrates the symmetric split for a reversal task with two available actions per problem; repeated-action problems such as (*a, a*) were excluded.

#### A.1.2 Episode structure

In the aggregate comparisons, episodes began with a presentation phase in which the available action identities (*a, b, c*, …) were presented sequentially, followed by null stimuli for any remaining setup steps. For the remainder of the episode, the network received a null stimulus together with the previous action *a_t_*_−1_ and reward *r_t_*_−1_. Episode lengths included the setup phase and matched the training rollout lengths in these runs.

- Episode setup duration: 6
- Trial length: 1
- Episode and rollout length: 64–128 timesteps, including setup (task dependent)

#### A.1.3 Task parameters

##### Probabilistic reversal task structure

- Num. total actions: 50
- Episode and rollout length: 64 timesteps, including setup
- Reward probability: 0.7
- Block switch probability: 0.1667

These settings describe the aggregate reversal comparisons. A separate earlier reversal configuration used reward probability 0.85 and block switch probability 0.1, with simultaneous action presentation and the same six-step setup and 64-step episode length.

##### 4-reversal task structure

- Num. total actions: 10
- Episode and rollout length: 128 timesteps, including setup
- Reward probability: 1.0
- Geometric block parameter: 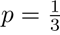
- Minimum block length: 3 trials.

Block lengths were sampled as *L* = *m* + *G* − 1, where *m* is the minimum duration and *G* ∼ Geometric(*p*) has support 1, 2,. … The switch hazard was zero before the minimum duration and *p* thereafter. No fixed maximum block length was imposed by this sampler; an episode could terminate before a sampled block was completed.

##### 3-arm drifting bandit task structure

- Num. total actions: 15
- Episode and rollout length: 128 timesteps, including setup
- Reward magnitudes: 1-9
- Gaussian drift kernel width: 0.15 of the reward range (*σ* = 1.2 reward units before discretisation and reflection)
- Reflection at bounds: True

##### Hierarchical reversal task structure

- Num. total actions: 10
- Episode and rollout length: 128 timesteps, including setup
- Reward probability: 1.0
- Lower-level geometric block parameter: *p*_lower_ = 0.45
- Upper-level geometric block parameter: *p*_upper_ = 0.4
- Minimum lower-level block length: 3 trials
- Minimum upper-level block length: 3 lower-level block completions

Both levels used the shifted geometric distribution defined above, without a fixed maximum duration. The lower-level counter advanced on each completed trial. The upper-level counter advanced only when a lower-level block ended, so upper-level changes occurred at lower-level block boundaries.

#### A.1.4 Bandit value estimation

For the bandit task we used a discretised Gaussian diffusion kernel for reward transitions between trials, with nominal width *σ* = 0.15(9 − 1) = 1.2 reward units and reflection at the boundaries [1, 9] (Figure 7a). Stacking the transition probabilities for each possible starting reward state gives a matrix *T*, whose rows index current states and columns index subsequent states. The expected reward after *n* transitions is given by *T ^n^***r**, where **r** = (1*, …,* 9)^T^ (Figure 7b). For each available action, we indexed this vector using the reward state last observed for that action, with *n* equal to the number of intervening drift transitions. All reward states drifted between trials, including the state of the just-sampled action; at the next decision, its estimate therefore included one transition since the observed reward. With increasing *n*, expected rewards converge to 5 regardless of the initial state (Figure 7d). This is the mean of the reflected process’s stationary distribution, which assigns probability 1*/*16 to each endpoint and 1*/*8 to each interior reward state.

**Figure 7:**
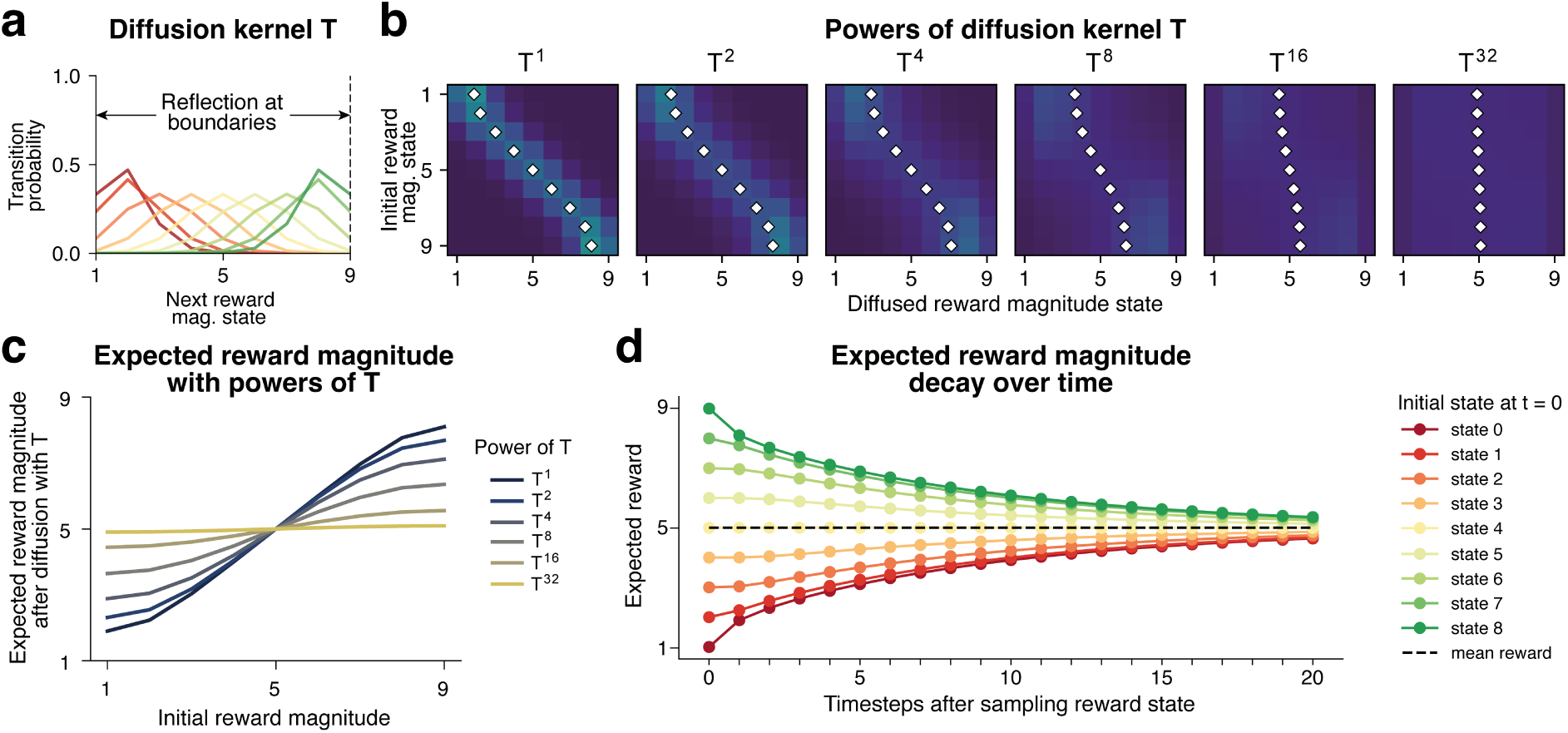
Expected value estimation of unsampled actions in the 3-arm magnitude bandit task. **a)** Diffusion kernel *T* for reward drift with reflection at boundaries. Colour denotes current reward magnitude; curves show the distribution of next reward magnitude. Colour code as in **d**. **b)** Diffusion kernel *T* visualised as a transition matrix and raised to successive powers, equivalent to the reward distribution after a specified number of diffusion steps. White diamonds denote the expected reward magnitude of each row (i.e., for each initial state). **c)** White diamonds from each plot in **b** visualised as curves. As *n* increases, expected reward approaches 5 for every initial reward state. **d)** Expected reward as a function of time since sampling. These curves were used to estimate expected values while the agent was not sampling the corresponding actions.

### A.2 Learning algorithm implementation

#### A.2.1 Reinforcement learning

##### Learning algorithm

Meta-RL networks were trained with PPO. Generalised advantage estimation (GAE) and value bootstrapping were truncated at trial boundaries. Recurrent hidden states were maintained across trials within each episode and reset at episode boundaries. Thus, truncation of the return targets did not reset the network’s recurrent memory between trials. Truncation simply allowed direct comparison of critic value output to the trial-by-trial value estimates of the ideal handcrafted agents.

A batch of rollouts was divided into 4 minibatches by randomly permuting the parallel environment sequences while preserving temporal order within each sequence. The minibatches were processed sequentially, updating the same network parameters. Each rollout batch was reused for 4 optimisation epochs, with a new permutation in each epoch.

A permutation-invariant auxiliary action-identification loss (Hungarian loss) encouraged retention of the available action identities after they were presented in the setup phase. For a problem with *K* available actions, the actor had *K* linear readout slots, each predicting an identity from the full action vocabulary. The loss at each timestep was

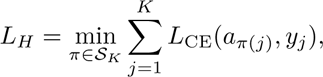

where S*_K_* is the set of permutations of the *K* available action identities, *y_j_* is the logit vector for slot *j*, and *L*_CE_ is cross-entropy. The implementation enumerated all *K*! assignments and selected the minimum summed loss. For two actions this reduces to

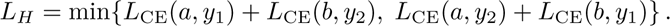

The loss was averaged across rollout timepoints and parallel environments, including presentation steps. It was computed from the actor readout and updated both the actor and shared input embeddings. The saved PPO runs used coefficient 0.1 for probabilistic reversal, cyclic reversal and magnitude bandit, and omitted the loss for hierarchical reversal. The supervised runs did not use this auxiliary objective.

##### Training parameters

- Parallel environments per rollout batch: 512 – 1024
- Max grad norm: 0.5
- Learning rate: 3e-4 – 1e-3
- Optimiser: AdamW (*ɛ* = 10^−5^)
- Weight decay: 0, except 3 x 10^−4^ for the saved reversal PPO runs
- Epochs per update: 4
- Minibatches per epoch: 4
- Discount factor *γ*: 1.0
- *λ_GAE_*: 1.0
- PPO clipping *ɛ_clip_*: 0.2
- Entropy loss coef: 0.001 – 0.01
- Value loss coef: 0.5
- Hungarian loss coef: 0.1 for reversal, cyclic reversal and magnitude bandit; 0 for hierarchical reversal

PPO networks combined 1) an input representation formed by summing learned embeddings of the stimulus, previous action and previous reward, 2) separate actor and critic RNNs receiving that shared representation, and 3) linear readout heads for the actor policy, critic value and auxiliary action-identity predictions where the Hungarian loss was enabled. The supervised architecture is described separately below. Gradients were clipped to a global norm of 0.5 before each optimiser update. Weight decay was applied to weight matrices, excluding biases and layer-normalisation scales.

##### Actor-critic network parameters

- Embedding dim: 128-256
- Actor hidden dim: 128-256
- Critic hidden dim: 128-256
- Cell type: GRU
- GRU candidate activation: ReLU
- Layer norm: Applied before the candidate activation
- Embedding weight init: Orthogonal(0.01)
- Input-to-hidden weight init: LeCun normal
- Recurrent weight init: Identity plus independent Gaussian noise with standard deviation 10^−3^
- Bias init: Zero
- Readout heads: Orthogonal(0.01)

#### A.2.2 Supervised learning

Supervised networks used the same input-embedding construction as the PPO networks, followed by a single recurrent network and a linear policy head. They had no critic or auxiliary action-identity head. During training, the network sampled and executed its own actions. A reference agent maintained its state using the executed action–outcome history and supplied an action target at each choice step. Networks were trained with cross-entropy loss on these targets at non-presentation choice steps, with one gradient update per collected rollout. This procedure trained the policy on the histories generated by its own behaviour.

The reference agents for cyclic and hierarchical reversal followed win-stay-lose-switch rules and were given the task’s action ordering or grouping. In cyclic reversal, an unrewarded choice prompted a switch to the next action in the cycle. In hierarchical reversal, the agent retained a remembered action for each set. After a first unrewarded choice it tried the other action in the current set; after consecutive losses exhausted that set, it switched to the next set and selected its remembered action. These rules changed the agent’s choices; block transitions in the environment followed the task dynamics independently.

For probabilistic reversal, a fixed-hazard reference observer used the task’s reward and reversal probabilities, initialised its belief that action A was the high-reward action to 0.5, and selected the action favoured by its current belief, breaking ties randomly.

For the drifting magnitude bandit, the reference agent selected the action with the largest expected immediate reward, breaking ties randomly. Expected rewards were calculated from the last observed reward state and the number of intervening drift transitions using the transition matrix described in Section A.1.4. Before all three actions had been sampled, the agent selected uniformly among the remaining unsampled actions. This was a greedy expected-reward policy; it did not explicitly optimise the future value of information.

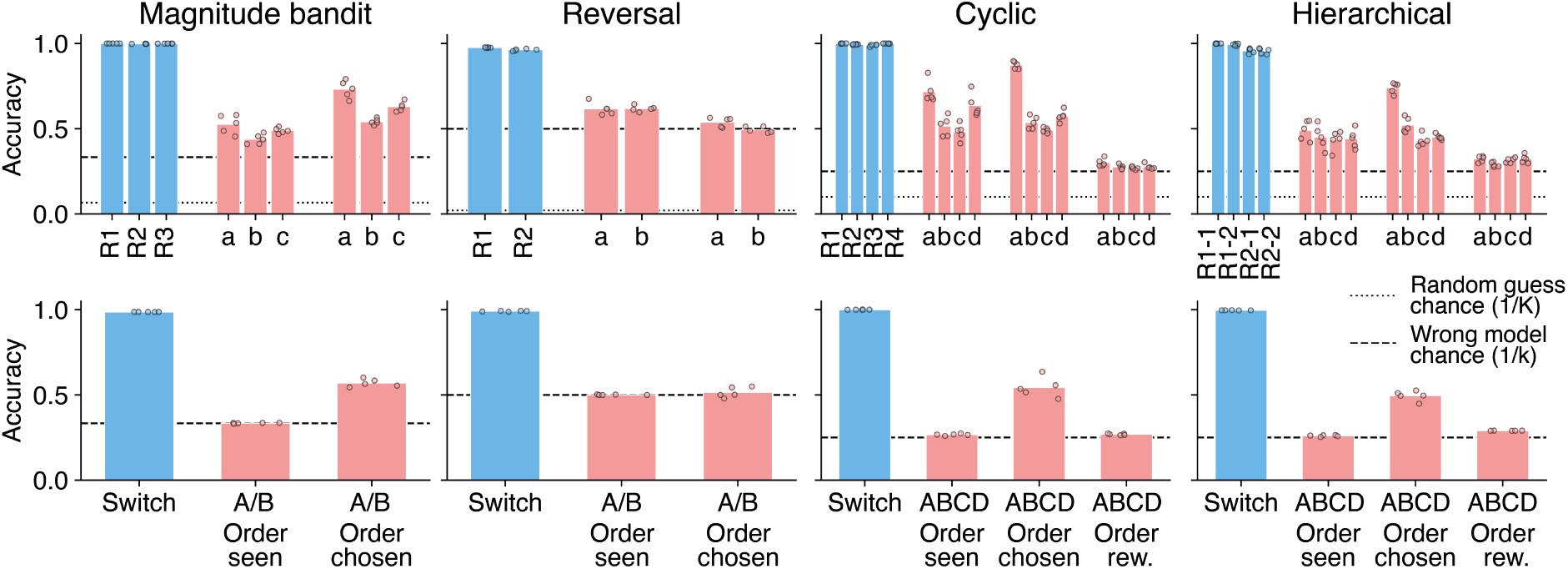

#### A.2.3 Network generalisation

We assessed generalisation by comparing performance on training problems and held-out combinations of the same action identities (Figure 9). We compared episode-level performance, performance conditional on the identity of the currently best action, and the frequency of invalid choices. These measures assessed transfer to the sampled held-out problems; they did not by themselves identify the representations responsible for that transfer. Behavioural generalisation was evaluated before applying the episode exclusions used for decoding.

**Figure 9:**
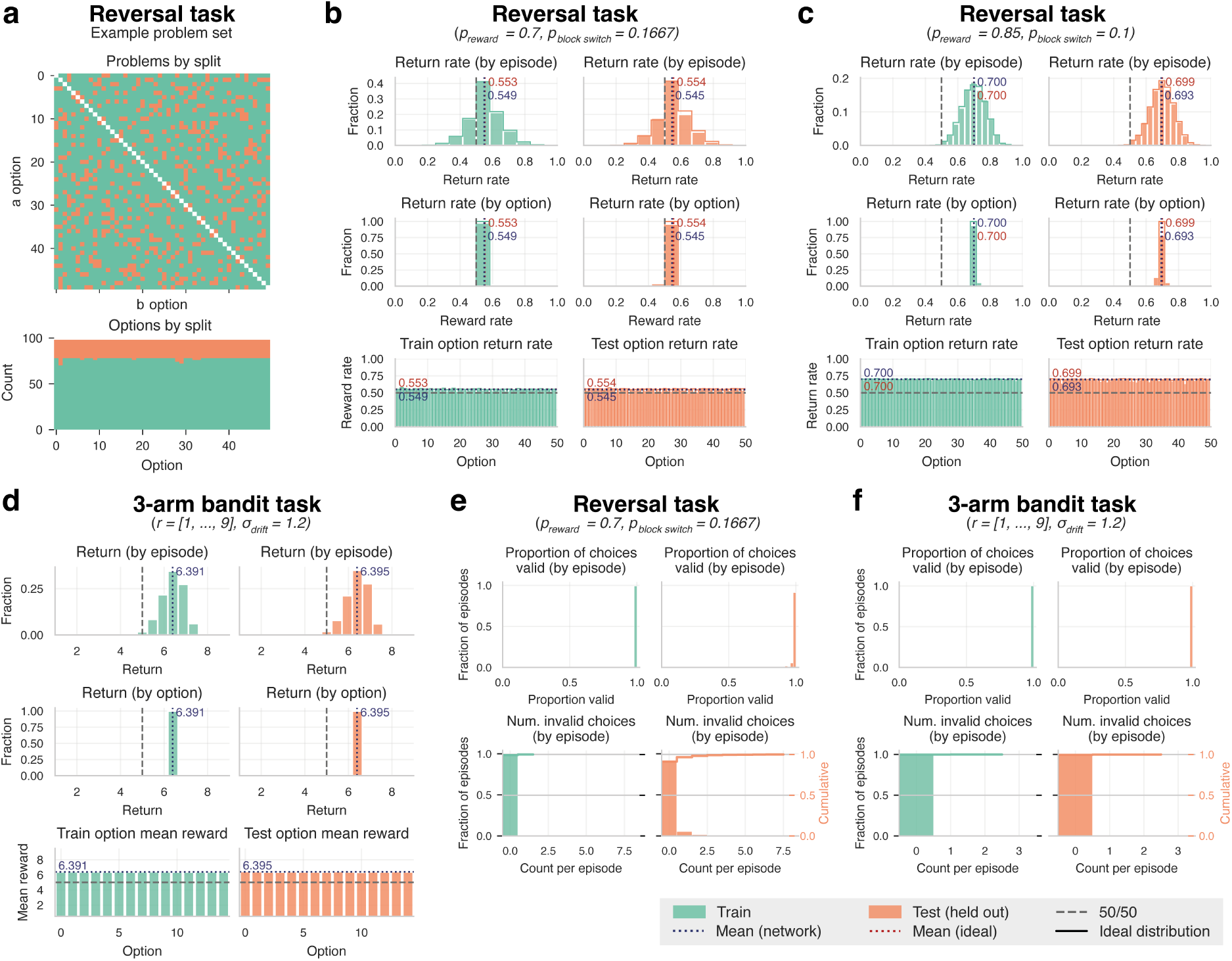
Generalisation to held-out problems in the reversal and 3-arm bandit tasks. **a)** Example train–test split. Problems comprise distinct action identities drawn from the task vocabulary (white diagonal, top). All permutations of each combination are assigned to the same split (symmetrical matrix, top), with approximately balanced coverage of individual identities (bottom). **b-d)** Performance on training and test problems, measured as reward rate in probabilistic reversal and mean reward per trial in magnitude bandit. Top row: distribution of episode-level reward rates or mean rewards per trial. Middle and bottom rows: distributions and averages conditional on each action identity being the environment’s currently best action. **e-f)** Choice validity. Invalid choices are actions outside the available action set and yield zero reward. Top row: distribution of the fraction of valid choices per episode. Bottom row: distribution of the number of invalid choices per episode.

#### A.2.4 Training and analysis replication

The saved working-memory and switching comparisons comprised five checkpoint runs for each combination of task and training regime (PPO or teacher-supervised learning). Decoder repetitions were first averaged within each checkpoint; plotted means and standard errors were then computed across checkpoints, with 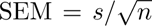, where *s* is the sample standard deviation and *n* is the number of checkpoints. A single-checkpoint analysis did not receive an across-checkpoint error bar.

For this collection, the configured training budget was 5 x 10^9^ environment timesteps, except for magnitude-bandit PPO seed 30, whose budget was 10^10^. Training configurations specified 400 evaluation occasions and greedy policy evaluation, with 1,024 episodes per problem split at each occasion; magnitude-bandit PPO seed 30 used 8,192. These budgets were configured limits, not necessarily the training duration attained by the analysed checkpoints. The configurations selected the latest saved checkpoint rather than the checkpoint with the highest evaluation return. Analysis loading also restored the latest available timestep when no timestep was specified.

Value decoding, encoding, geometry and causal analyses used the individual checkpoints listed in Table **??**. Post-training analysis simulations were distinct from the evaluation batches collected during training. They generally targeted 10,000 episodes per problem split, rounded down to a whole multiple of the number of problem layouts; eligibility and exclusion criteria were then applied as described for each analysis.

### A.3 Analyses

#### A.3.1 Decoding

For the baseline decoding analyses, the decoders were trained on episodes from the network training problem-set and tested on episodes from the network test problem-set. Therefore, all reported baseline decoding results are on problems that the network itself was not trained on. For decoder training, the network train problem-set was divided into train and validation sets. The split was performed in such a way that all permutations of a problem (e.g. (*a, b*) and (*b, a*)) were in the same split, and that each individual action was approximately represented in the specified train/val ratio.

Baseline decoding accuracies were evaluated on the network *test* problem set. High accuracy therefore indicates that the decoded information generalises to held-out problems. Decoder readouts following causal interventions were evaluated separately on manipulated states, as described in Section A.3.4.

For decoding, logistic regression was used with the sklearn package. Balanced class weights were used, particularly for value decoding, where the class distribution was highly uneven. Hidden-unit activity was generally standardised using a StandardScaler fitted to the decoder-training observations and then applied to validation and test observations. Unless otherwise specified, the inverse regularisation strength was *C* = 1; the saved bandit baseline value decoders used *C* = 0.01.

Working memory decoders used non-presentation choice steps. For probabilistic reversal, eligible steps began at trial index 0, including the first choice. For cyclic and hierarchical reversal, every available action had to have delivered a positive reward at least once; the timestep on which the last required reward was delivered was included. For the bandit baseline analyses, eligible steps had trial index at least 40 and all three actions had to have been sampled. Observations with missing targets were excluded separately for each decoder, so the retained observations could differ between targets.

For reversal baseline value decoding, targets were the reference observer’s belief *p_A_* that action A was the high-reward action, and its folded confidence *p*_max_ = max(*p_A_,* 1 − *p_A_*). The latter describes confidence in the observer’s preferred action, which need not coincide with the action selected by the RNN. Complementarity of the observer’s two beliefs does not constrain the number of value representations in the RNN. The critic’s scalar output was also decodable from the actor RNN activity, and was used for the reversal value-manipulation analysis (Section A.3.4). For the 3-arm bandit task, action values were estimated from the reward received when each action was last sampled and the number of transitions since that observation. We propagated the observed reward state through the task’s drift transition model and took the expected reward under the resulting distribution (Section A.1.4). The estimated values were assigned to behavioural ranks or sorted numerically, as specified for each analysis below.

For reversal decoding, entire episodes were excluded if any choice was outside the available action set. Exclusion was applied separately to training and test episodes. In the inspected reversal baseline run, approximately 0.1% of training episodes and 12.4% of test episodes were excluded by this criterion. The presentation phase was excluded from all baseline decoding analyses.

Continuous value-related targets were discretised into equally spaced bins. The saved reversal baseline analysis used 20 bins; the 3-arm bandit baseline analysis used 10 bins. Percentile limits defined the binning range, and observations beyond that range were assigned to endpoint bins rather than excluded. The reversal belief binning used eligible training and test observations: for *p_A_*, the limits were symmetric about 0.5 and determined by the larger of their 99th percentiles; the folded-confidence analysis used a separately constructed range or the same edges as *p_A_*, as specified by the comparison. Binning for the causal analyses is described separately below.

#### A.3.2 CPD and encoding analyses

We used linear encoding models to compare ranked working memory and value codes with action-specific value codes in the 3-arm bandit task. Encoding predicts network activity from task variables, allowing predictor sets of different sizes to be compared against the same hidden activity.

We used actor RNN hidden activity at each choice, after processing the current observation and before receiving the outcome of that choice. Episodes were generated for the network training and test problem sets. Training problems were further divided into encoder training and validation sets in an approximately 0.8:0.2 ratio, keeping all permutations and repetitions of a problem together. Prediction performance was evaluated on the network test problem set. We included non-presentation choice steps from trial index 40 onwards, after all three actions had been sampled, and retained only observations with a complete inferred ranking and finite value estimates. All compared encoding models used the same observations.

Working memory ranks were inferred retrospectively from runs of repeated choices among the available actions. Rank 1 was the action selected in the current run, rank 2 was the action selected in the next distinct run, and rank 3 was the remaining action. The final run of an episode was excluded because its subsequent choice was unavailable. Expected values were computed from the sampling history using the transition model described in Section A.1.4, and assigned to the corresponding behavioural rank slots. This preserved the association between each action identity and its estimated value.

The full model contained three candidate predictor blocks. Ranked working memory comprised three one-hot encodings of action identity, one for each rank, giving 3 x 15 predictors. Ranked value comprised three one-hot encodings of discretised expected value, giving 3 x *B* predictors for *B* value bins. The Q-value representation instead assigned value-bin indicators to fixed physical action identities, giving 15 x *B* predictors. Only the three actions available in the current problem had active value indicators; unavailable actions contributed zeros. The Q-value block therefore represented the same observer estimates in action coordinates, rather than values obtained by fitting a separate temporal-difference learning model.

We included two nuisance predictor blocks in every model. The first supplied additive predictors for action presence: a 15-dimensional binary vector with three active entries indicating the available identities. The second was a one-hot encoding of the preceding action–outcome pair (*a_t_*_−1_*, r_t_*_−1_), with 15 x 10 possible categories for the 15 action identities and reward inputs 0–9. This allowed a separate fitted effect for each pair, including action–reward interactions. Both blocks entered the joint encoding model to account for activity associated with the available actions and preceding action/reward input.

Predictor columns were centred, and hidden units were centred and standardised, using encoder-training statistics. The same transformations were applied to validation and test data. A linear encoding model was fitted for each subset of the candidate blocks:

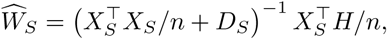

where *X_S_* contains the centred nuisance predictors and candidate blocks in subset *S*, *H* is standardised hidden activity, *n* is the number of training observations, and *D_S_* is the diagonal regularisation matrix. A nuisance-only model provided the baseline prediction. For each full or reduced model, all retained coefficients, including nuisance coefficients, were fitted anew on the same observations.

For each hidden unit *j*, held-out prediction performance was measured as

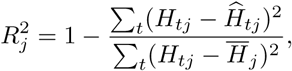

where *t* indexes the retained choice observations across episodes and 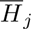 is the mean within the evaluation set. We averaged 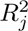 across hidden units to obtain 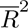, excluding units whose evaluation-set total sum of squares was below 10^−12^. The coefficient of partial determination (CPD) for predictor block *k* was

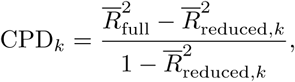

where the reduced model omitted block *k* and retained both nuisance blocks and all other candidate blocks. The denominator was bounded below by 10^−12^ for numerical stability. CPD measured the improvement in held-out prediction attributable to that block, conditional on the retained predictors, relative to the variance remaining unexplained by the reduced model. Reported CPDs were expressed as percentages; negative values indicated that including a block worsened held-out prediction. Because the candidate representations are correlated, a small CPD does not imply that the corresponding information is absent from network activity.

#### A.3.3 Subspace angles

For the 3-arm bandit task, we fitted six logistic regression decoders to actor RNN activity: three for action identities occupying the behavioural ranks and three for discretised observer values sorted in descending order. Behavioural ranks were inferred from choice blocks as described above; their ordering could differ from the ordering of observer-estimated values.

All six decoders used the same eligibility and sampling masks and decoder train/validation split. Observations were restricted to non-presentation choice steps from trial index 40 onwards, after all three actions had been sampled. Observations without complete ranked targets were excluded. Classes were weighted inversely to their frequency during fitting, and the inverse regularisation strength was *C* = 0.01 for these geometry decoders.

For each decoder, we obtained a coefficient vector **w***_c_* for each fitted class *c*. We removed the mean coefficient vector across classes,

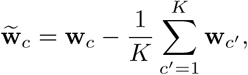

where *K* is the number of fitted classes. This removes the coefficient component shared by all class logits.

For each working memory rank *r* and value rank *s*, we calculated the acute angle between every action-identity coefficient vector and every value-bin coefficient vector:

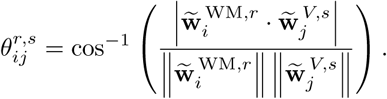

Angles were expressed in degrees. Taking the absolute dot product treats parallel and antiparallel vectors as the same axis. Zero-norm vectors were excluded. The 3 × 3 heatmap reports the mean angle across all valid action-identity/value-bin pairs for each rank pairing. Values approaching 90^◦^ indicate approximately orthogonal class-contrast directions.

We also calculated principal angles between the complete decoder subspaces. For each class-centred coefficient matrix, singular value decomposition provided an orthonormal basis *Q* for its row space. Singular directions were retained when their singular value exceeded 10^−8^ times the largest singular value. For a pair of bases, the principal angles were

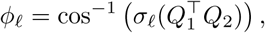

where *σ_ℓ_* denotes a singular value. Principal angles describe the orientation of the decoder spans; the heatmap reports the mean angles between individual class-contrast vectors.

#### A.3.4 Causal manipulations

We tested the effects of replacing working memory identities and value representations on subsequent choices and working memory contents in the reversal and 3-arm bandit tasks. For each manipulated variable *x*, we fitted a linear encoding model

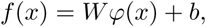

where *φ*(*x*) is a one-hot encoding of its identity or value category. The model was fitted by ordinary least squares to actor RNN hidden activity using the encoder-training problems. We then subtracted the predicted activity for the original variable and added the prediction for a target variable:

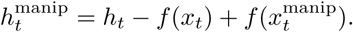

This replaced the encoder-predicted activity pattern while retaining the residual. The fitted pattern could include activity shared with correlated task variables, so the intervention was not assumed to isolate uniquely attributable variance.

Within the recurrent network, only the actor GRU hidden state was changed; the critic hidden state was retained. The modified state was inserted after action selection and before processing the following action/reward observation. Simulations resumed using the saved environment state, observer state and unchanged observation from the next timestep. Paired control and manipulation conditions used the same random keys. Interventions and associated decoder readouts used states sampled from simulated network-training problems.

##### Working memory identity replacement

For the reversal task, we replaced the identity of the chosen or unchosen action. For the bandit task, we replaced the identity occupying each of the three behaviourally inferred ranks in separate manipulations. Each target identity *i* was mapped to a different identity according to

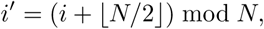

where *N* is the full number of action identities. Events were excluded if the mapped identity duplicated another action in the problem.

We also replaced the corresponding available action in the environment, preserving its position and associated reward process. The injected identity therefore inherited the reward association of the original action. In the reversal task, the identity assigned to the rewarded action was updated accordingly. Actions were not presented again, and the next observation was unchanged. The paired control retained the original hidden state and environment identities.

Identity encoders were fitted to non-presentation choice steps from trial index 20 onwards. For the bandit task, the first 40 simulation timesteps were also excluded from encoder fitting. Candidate intervention events were sampled with probability 0.1 from reward steps at trial index 10 or later, subject to sufficient remaining episode duration. The immediately preceding hidden state was replaced, and paired trajectories continued for 20 steps in the reversal task and 50 steps in the bandit task.

At each elapsed step *τ*, we measured the probability of selecting the injected identity conditional on the control selecting the original target:

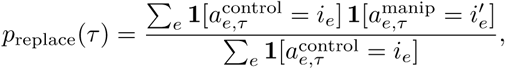

where *e* indexes intervention events. The conditioning was applied independently at each elapsed step. Corresponding probabilities for the original identities used the same denominator. The curves therefore measured substitution when the paired control selected the targeted action. Reward-matched trajectories were selected for illustration; aggregate curves used the full eligible event set (Figure 10).

**Figure 10:**
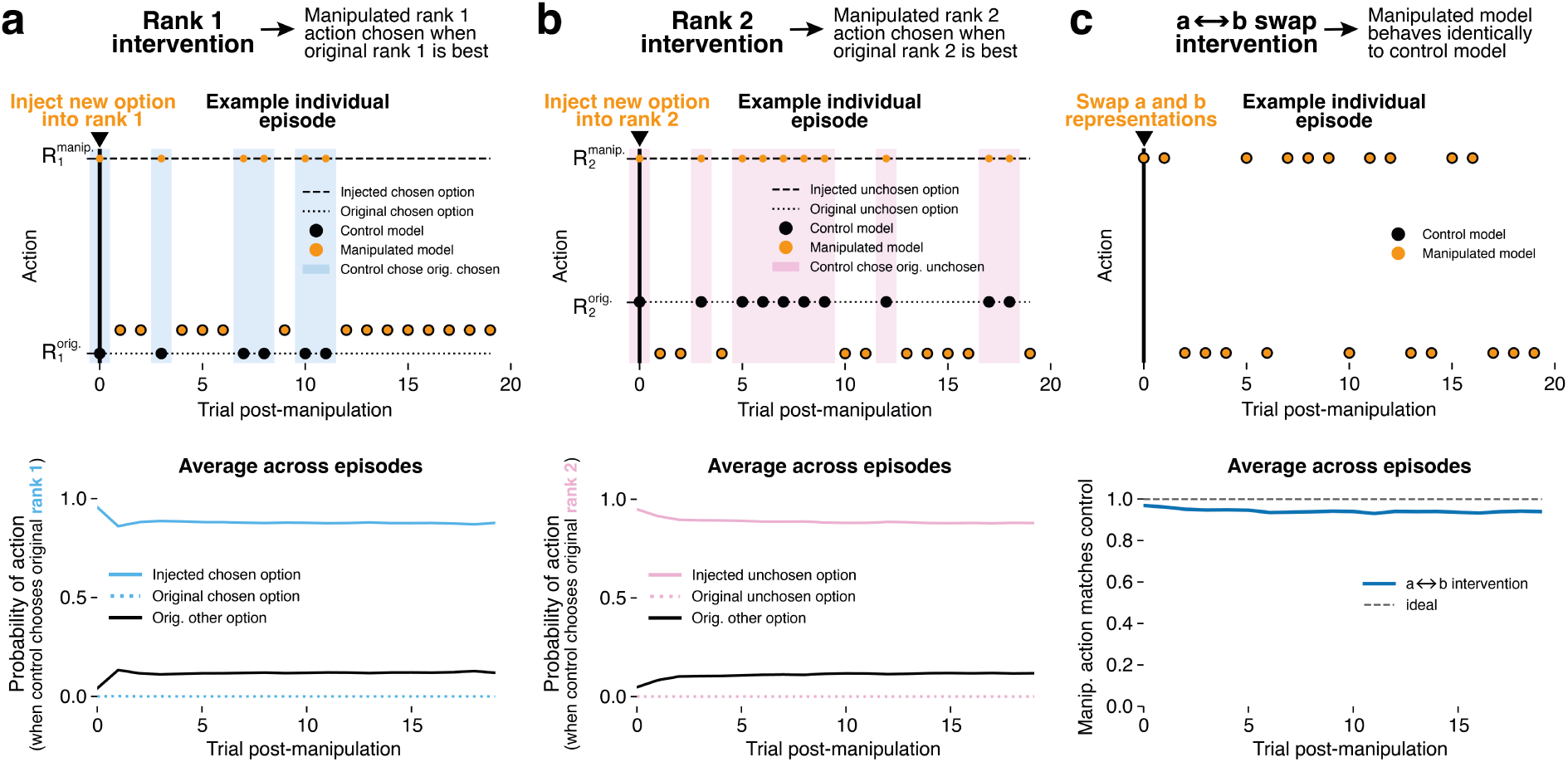
Causal manipulations of ranked action subspaces in reversal task predictably affects future behaviour. **a,b)** Top: behaviour in example episode following manipulation. For each rank manipulation, when target rank is best (shaded pastel regions) the manipulated RNN chooses the injected option rather than original option corresponding to that rank. Bottom: in shaded regions only (original target rank becomes best) across all episodes, probability of manipulated RNN choosing injected option is high. **c)** Swapping the contents of the static a and b subspaces that we found were decodable in reversal task (Figure 3f left) has no effect on behaviour. Top shows identical actions taken as un-manipulated model on an example episode, bottom shows the average match across all episodes.

##### Value replacement in the reversal task

For reversal, the critic’s scalar value output was used as the value predictor for actor activity. Values were divided into 20 equally spaced bins, with the lower limit given by the smaller of the masked training and test first percentiles, and the upper limit by the larger of their 99th percentiles. Values beyond these limits were assigned to the endpoint bins. The encoder was fitted to non-presentation choice steps from trial index 1 onwards.

We selected unrewarded transitions at the reward-delivery step, with sufficient remaining episode duration, replaced the original fitted value pattern with the pattern for each target bin in turn, and simulated one further timestep. Every target bin was tested on the same selected states, with an unmodified paired control. Environment identities, reward dynamics and critic hidden states were unchanged. We measured the fraction of next choices matching the previously chosen action (stay) or previously unchosen action (switch), both overall and as a function of the original value bin. Chosen- and unchosen-action decoders were applied to the actor state after the subsequent network update, and we measured the fraction of predictions matching each original identity (Figure 11).

**Figure 11:**
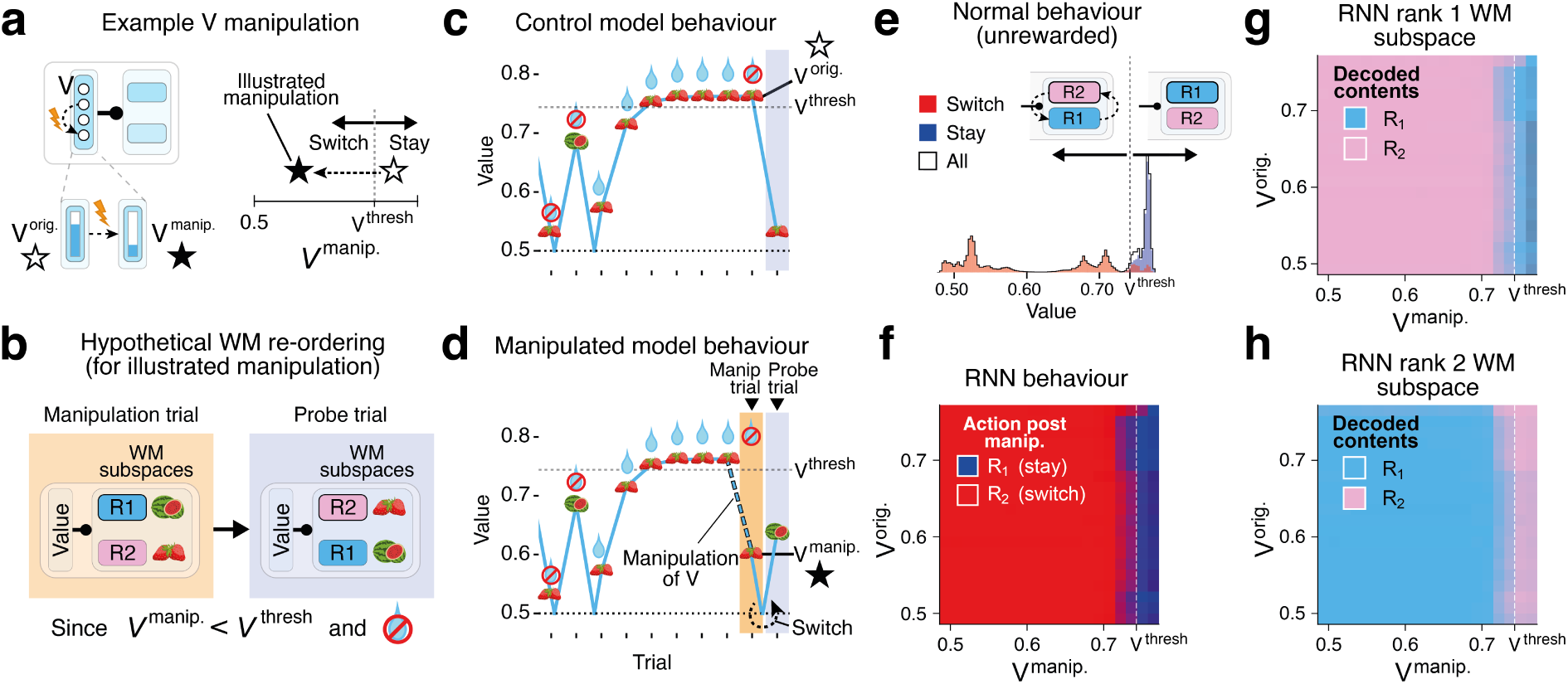
In the reversal task, manipulation of value of (actor) RNN predictably affects behaviour and subspace contents. **a)** Left: schematic of manipulation targeting value subspaces. Curved dotted arrow between neurons in the model denotes change from original value to new manipulated value. Right: Solid star and empty star denote pre- and post-intervention value respectively for the illustrated manipulation on the left. *V ^thresh^* denotes the behavioural threshold below which an unrewarded trial triggers a switch. **b)** Illustrated value manipulation triggers hypothetical re-ordering of contents of WM subspaces on subsequent trial. R1/2 (blue/pink) i.e. chosen/unchosen denote the option stored in rank 1/2 WM subspaces just prior to manipulation. **c)** Hypothetical behaviour of control (un-manipulated) model when unrewarded but above *V ^thresh^*. Value decreases but does not trigger a switch. **d)** Hypothetical behaviour of manipulated model when unrewarded but below *V ^thresh^*. Value decreases enough to trigger a switch — R2 (watermelon) becomes more valuable than R1 (strawberry) so is selected post-intervention. **e)** Distribution of switch and stay decisions by the network with respect to value (unrewarded trials only). **f)** Manipulation predictably affects RNN behaviour: on unrewarded trials, when *V ^manip^ < V ^thresh^*, R2 (red) is chosen on the post-intervention trial, otherwise R1 (blue) is chosen. This effect is independent of the original value prior to manipulation, *V ^orig^*. **g,h)** Memories are moved between ranked subspaces when manipulating ranked value across the critical threshold *V ^thresh^*.

##### Value replacement in the 3-arm bandit task

For the bandit task, we manipulated the second- and third-largest observer-estimated values jointly. These value ranks were obtained by sorting the estimates numerically; behavioural rank labels were retained when assessing the identities represented by the working memory decoders. Each value was divided into 15 equally spaced bins with common limits. The lower limit was the larger of the minimum observed values for ranks 2 and 3, and the upper limit was the midpoint of the reward range, 5, with a numerical tolerance of 10^−6^. A joint category identified each pair of bins, allowing the encoder to fit a separate activity pattern for each value combination represented in its fitting data.

We included non-presentation choice steps from trial index 40 onwards, with complete behavioural rank labels, both original value estimates strictly within the binning limits, and at least one subsequent timestep available. For each joint category observed in the eligible baseline data, the encoder’s prediction replaced the original joint value pattern in every selected state. The environment and critic state were unchanged, and simulations continued for one timestep using the same initial states and random keys across target conditions.

Analyses were performed separately for transitions delivering rewards of 2, 3 or 4. We measured the fractions of subsequent actions matching the original behavioural rank-1, rank-2 and rank-3 identities. We also applied all three working memory decoders immediately after replacement and after the subsequent network update, measuring how often each decoder reported each original identity. To combine reward conditions, the two-dimensional value grids were centred on the bin containing the received reward *r_t_*, and averaged with equal weight across reward conditions wherever entries were available. At the resolution of the value bins, the resulting coordinates describe manipulated values relative to that reward, 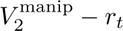 and 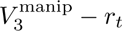. This reference should be distinguished from the observer’s highest estimated value, which need not equal the received reward.

## Footnotes

1 Note that in the original Harlow task, value is indicated by deterministic reward and therefore does not need to be integrated over time. In this special case, reward input can act directly as a control signal for swapping WM contents. In other tasks that do require value integration but have perfectly anti-correlated values (such as probabilistic reversal tasks where *V*_1_ a −*V*_2_), only one value representation is required rather than separate *V*_1_ and *V*_2_ representations.

